# eIF2B Selectively Anchors and Activates Mutant KRAS

**DOI:** 10.1101/2025.11.10.686860

**Authors:** Hyungdong Kim, Shiqi Diao, Kwang-Jin Cho, Hyun-Ro Lee, Junchen Liu, Pascal Egea, Tatu Pantsar, Milla Kurki, Nour Ghaddar, Shuo Wang, Jia Yi Zou, Mehdi Amiri, Ritchel Gannaban, John Hancock, Kylie M. Rice, Qiyun Deng, Atsuo Sasaki, John Asara, Brajendra Tripathi, Douglas Lowy, Rosalie Lawrence, Maria Hatzoglou, Carlos R Azpilcueta-Nicolas, Jean-Philip Lumb, John Columbus, Thomas J Turbyville, Christopher B. Marshall, Mitsuhiko Ikura, Jay T. Groves, Nahum Sonenberg, Peter Walter, Antonis E. Koromilas

## Abstract

Much is known about how RAS oncoproteins regulate mRNA translation factors, but the reverse relationship, how translation factors influence RAS activity, has remained largely unexplored. At the plasma membrane (PM), Son of Sevenless (SOS) acts as the canonical guanine nucleotide exchange factor (GEF) for RAS proteins, yet mechanisms governing its specificity for individual RAS isoforms remain unknown. Here, we show that the translation initiation factor eIF2B, best known for its GEF function in translation initiation, forms a distinct complex with SOS and mutant KRAS at the PM, but not with other oncogenic RAS variants. Mechanistically, eIF2B acts as an allosteric regulator of SOS, selectively enhancing GDP–GTP exchange on mutant KRAS. This specificity arises from the translational activity of eIF2B, which upregulates glycosphingolipid (GSL) biosynthesis to remodel PM lipids and preferentially anchor mutant KRAS. Together, our results uncover an unexpected moonlighting function of eIF2B: acting both as a direct activator of SOS and as a regulator of GSL pathway that shapes the membrane landscape, both required for mutant KRAS activation. These insights redefine our understanding of eIF2B and mutant KRAS functions in cancer and have profound implications for KRAS-driven oncogenesis.

**Graphical Abstract:** 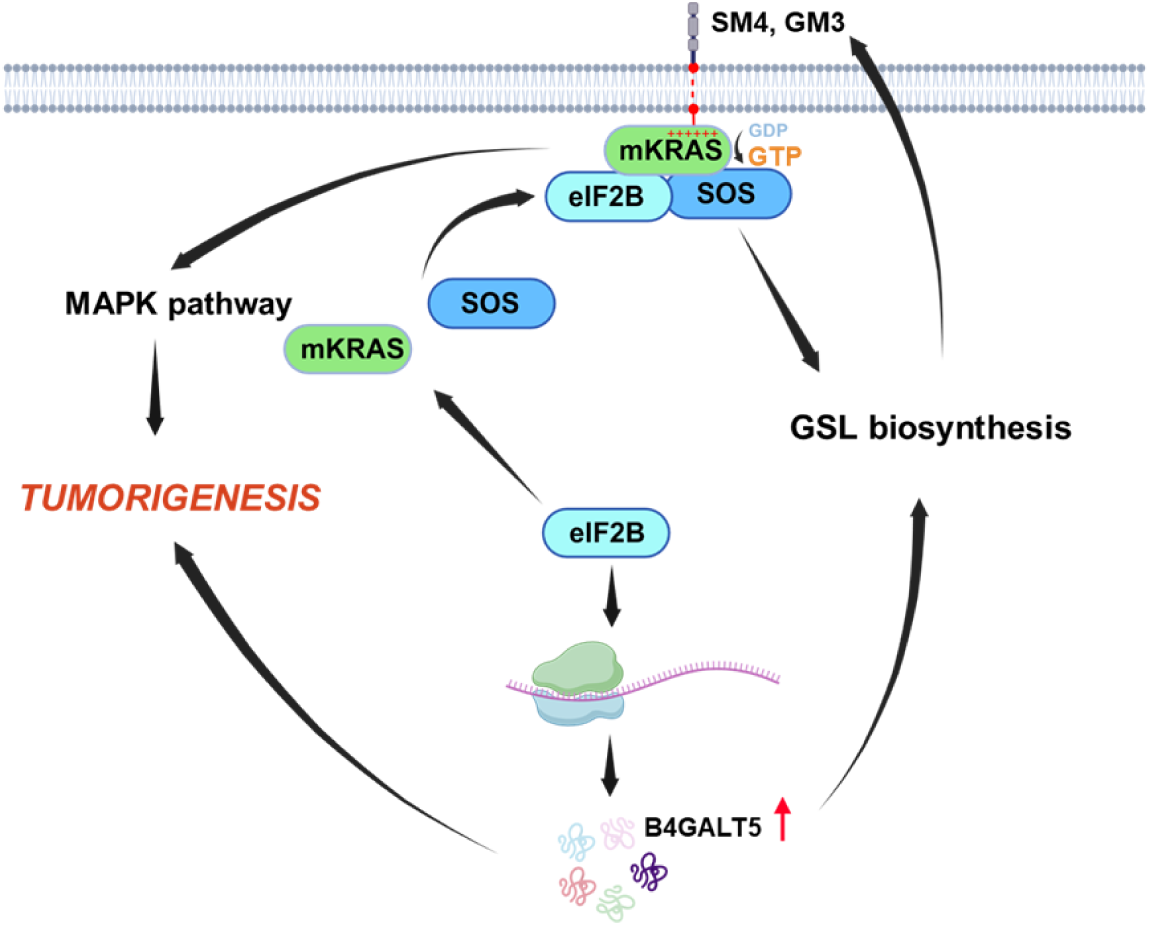

- eIF2B interacts with mutant KRAS and SOS at the plasma membrane (PM).
- The eIF2B:SOS complex promotes the GTP-bound active state of mutant KRAS.
- eIF2B enhances the translation of B4GALT5 mRNA, encoding a key enzyme of glycosphingolipid (GSL) biosynthesis.
- Upregulation of the GSL metabolites, ganglioside GM3 and sulfatide SM4, remodels PM lipid composition to facilitate eIF2B:SOS:KRAS complex formation and mutant KRAS nanoclustering.
- Through its interaction with SOS and stimulation of GSL synthesis, eIF2B selectively activates mutant KRAS at the PM among RAS isoforms.
- eIF2B is required for the development of mKRAS-driven lung adenocarcinoma in mice.
- eIF2B is a marker of poor prognosis in mutant KRAS-driven cancers.

## INTRODUCTION

The Kirsten Rat Sarcoma Virus (KRAS) protein is a small, membrane-bound protein (∼189 amino acids in length) that plays a crucial role as regulator of cell proliferation (1). KRAS possesses intrinsic GTPase activity, enabling it to cycle between an active GTP-bound state and an inactive GDP-bound state (1). This cycling is tightly regulated by GTPase-activating proteins (GAPs) and guanine nucleotide exchange factors (GEFs) (2).

Somatic gain-of-function mutations in *KRAS* are prevalent in three of the most lethal cancers worldwide: lung, pancreatic and colorectal cancers (3). These mutations typically involve single amino acid substitutions at positions G12, G13, or Q61, which favor the GTP-bound active state of KRAS (3). Despite the preferential GTP binding, most of the oncogenic mutant KRAS proteins retain intrinsic GTP hydrolysis activity, which can be stimulated by GAPs (4) and counteracted by GEFs (5, 6).

KRAS activation drives multiple intracellular signaling pathways, with the RAF-mitogen-activated protein kinase (MAPK) pathway, comprising MEK and ERK, being particularly prominent (7). Like other oncoproteins, mutant KRAS relies on mRNA translation to selectively synthesize proteins essential for tumor growth (8).

Recent therapeutic advances have led to development of mutant KRAS inhibitors targeting either the inactive GDP-bound state (KRAS^OFF^) or the active GTP-bound state (KRAS^ON^) (9–11). However, resistance remains a major challenge, emphasizing the need for a deeper mechanistic understanding of mutant KRAS function to guide the development of more effective therapies (12).

The translation initiation factor eIF2B is a crucial protein complex involved in eukaryotic protein synthesis. It plays a central role in regulating mRNA translation by catalyzing the exchange of GDP for GTP on the eukaryotic initiation factor eIF2 complex, a process essential for translation initiation (13). Only the GTP-bound form of eIF2 can participate in forming the translation initiation complex, which delivers the initiator tRNA to the ribosome. Under stress, eIF2 is phosphorylated on serine 51 of its alpha (α) subunit by various kinases (e.g., PERK, GCN2, PKR, HRI)(14), converting it into a competitive inhibitor of eIF2B (15). This inhibition of eIF2B, a defining feature of the integrated stress response (ISR), decreases global protein synthesis, conserving resources and helping the cell manage stress. Despite this general reduction in translation, specific mRNAs involved in stress responses remain actively translated. Some of these mRNAs encode transcription factors that drive transcriptional reprogramming, enabling cells to survive and adapt to stress (16).

eIF2B is a large (∼ 600 kDa) complex composed of five subunits (α, β, γ, δ, ε), forming a decameric structure with two pentamers arranged in mirror-image symmetry. eIF2B cycles between an active state, when it is bound to non-phosphorylated eIF2, and an inactive state when bound to phosphorylated eIF2 (15).

Previous studies have shown that dysregulated eIF2B expression contributes to tumorigenesis and influences therapeutic responses (17–19). However, it remains unclear to what extent these functions depend on its translational function through the regulation of its GEF activity by phosphorylated eIF2, especially given evidence that eIF2B can also operate independently of phosphorylated eIF2 (20).

Through cellular and mouse models, we identify eIF2B as an unexpected critical partner of mutant KRAS and a key driver of its oncogenic activity. eIF2B forms a plasma membrane (PM)-associated complex with SOS and mutant KRAS, thereby selectively stimulating SOS GEF activity for mutant KRAS over other RAS oncoproteins. The formation of the PM-localized complex is facilitated by eIF2B-dependent stimulation of glycosphingolipid (GSL) biosynthesis, which remodels the PM lipid landscape to preferentially support mutant KRAS anchoring. Genetic inactivation of eIF2B selectively suppresses mutant, but not wild-type, KRAS tumor growth, establishing eIF2B as a key molecular hub linking translational control to activation of oncogenic KRAS.

## RESULTS

### eIF2B activates the MAPK pathway and sustains mutant KRAS cell survival

To investigate the role of eIF2B in KRAS signaling, we silenced its catalytic ε subunit, required for its GEF activity toward eIF2 (15), using siRNAs and shRNAs in lung adenocarcinoma (LUAD) H358 and pancreatic carcinoma Mia-PaCa-2 cells, both harboring the *KRAS G12C* allele, as well as in LUAD H1703 cells containing wild-type *KRAS*. RNA sequencing (seq) followed by KEGG pathway enrichment analysis of differentially expressed genes in control and eIF2Bε knockdown (KD) cells revealed a marked suppression of the MAPK pathway specifically in mutant KRAS cells, but not in H1703 cells with wild-type *KRAS* (**Figure 1a**).

**Figure 1.**
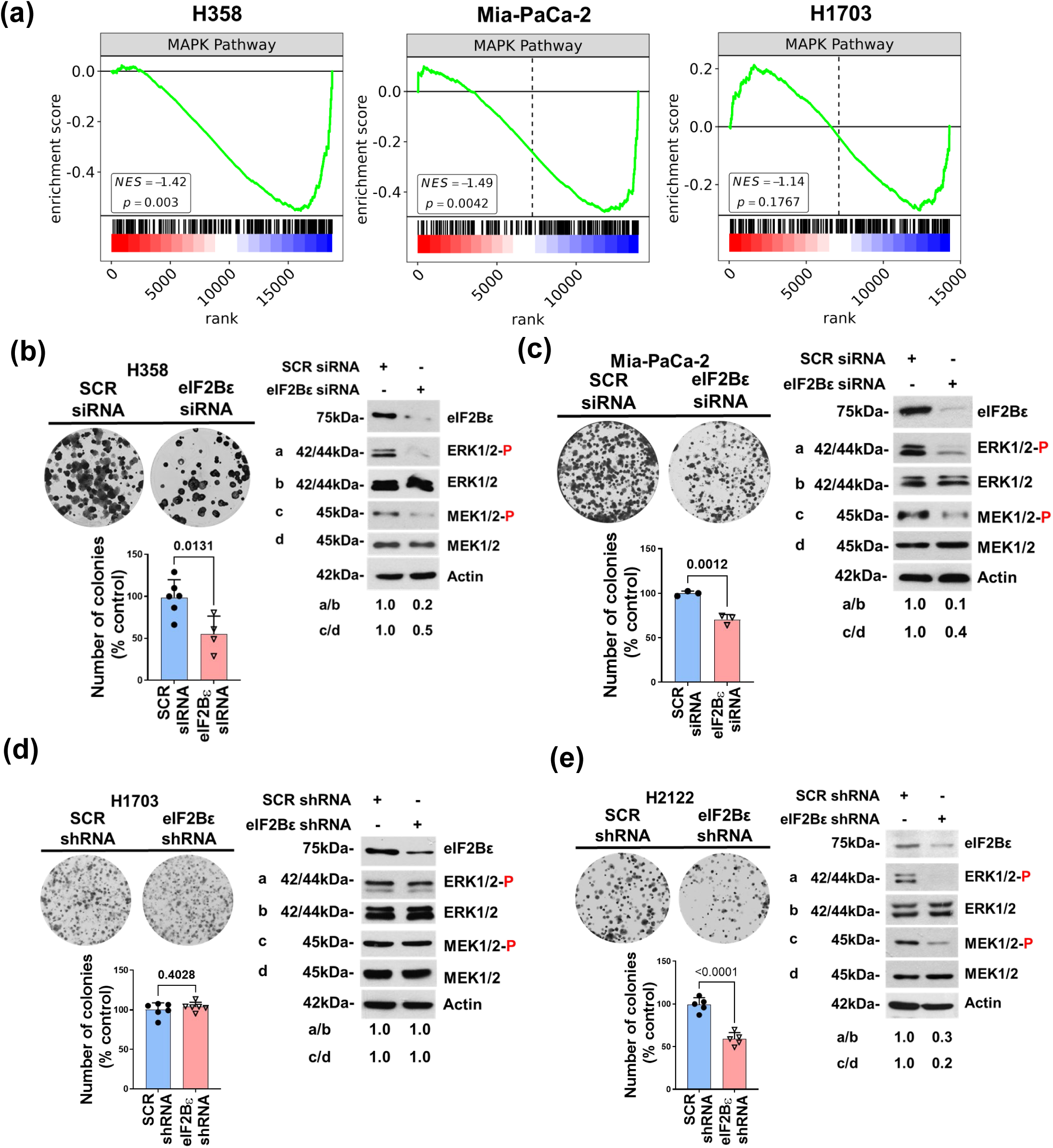
eIF2B supports survival and MAPK pathway activation in mutant KRAS cells. (**a**) KEGG pathway enrichment analysis of transcriptomic data from control and eIF2Bε KD H358 and MiaPaCa-2 cells harboring *KRAS G12C*, as well as H1703 cells with wild-type *KRAS*. The negative enrichment scores for the MAPK pathway following eIF2Bε KD in mutant but not wild type KRAS cells support a positive regulatory role of eIF2Bε in this signaling cascade. (**b–e**) Assessment of colony-forming ability and MAPK pathway activity in human cancer cell lines harboring either mutant KRAS (H358, MiaPaCa-2, H2122) or wild type KRAS (H1703) following eIF2Bε KD using siRNA or shRNA. SCR, scrambled siRNA or shRNA used as a negative control. Protein lysates were analyzed by immunoblotting for phosphorylated and total MEK and ERK. The ratio of phosphorylated to total protein is indicated. Graphs show quantification from three independent biological replicates; each performed in triplicate. Data are presented as mean ± SEM.

Consistent with this, eIF2Bε KD by either siRNAs or shRNAs resulted in 50-80% reduction of MEK and ERK phosphorylation and impaired colony formation in H358 and Mia-PaCa-2 cells, while having no effect on H1703 cells (**Figure 1b-d**, and **Suppl Figure 1a**). Similarly, in LUAD H2122 cells, which carry two *KRAS G12C* alleles, eIF2Bε KD also suppressed MAPK signaling and reduced cell survival (**Figure 1e**).

To further assess eIF2B’s role in mutant versus wild type KRAS-driven signaling, we used H1703 cells, which endogenously express wild-type KRAS, and engineered them to express green fluorescent protein (GFP)-tagged KRAS G12C or wild-type KRAS (**Suppl Figure 1b**) (21). Expression of GFP-KRAS G12C sensitized H1703 cells to eIF2Bε KD, resulting in suppressed MAPK signaling and reduced colony formation. This effect was not observed in cells expressing GFP-wild-type KRAS. Moreover, we tested colorectal adenocarcinoma HCT116 cells harboring *KRAS G13D* and their isogenic counterpart, HK2-8 cells, which express wild-type *KRAS* (22) (**Suppl Figure 1c**). Consistently, eIF2Bε KD impaired MAPK signaling and cell survival in HCT116 cells, but not in HK2-8 cells, indicating that the dependency on eIF2B is specific to mutant KRAS-expressing cells.

We also investigated the effects eIF2Bε silencing in mouse LUAD cells harboring *KRAS G12D*, either proficient (eIF2α^S/S^) or deficient in phosphorylation of eIF2α at serine 51 (eIF2α^A/A^), established from an autochthonous LUAD model (21). Since phosphorylated eIF2α antagonizes eIF2B’s GEF activity and translational output, we aimed to examine whether eIF2B’s ability to promote cell survival and MAPK signaling is regulated by phosphorylated eIF2α. Colony formation assays showed that eIF2α^A/A^ cells had reduced survival compared to eIF2α^S/S^ cells, consistent with previous findings that phosphorylated eIF2α promotes cell survival in KRAS-driven lung cancer (21) (**Suppl Figure 1d**). However, eIF2Bε KD impaired the survival and MAPK signaling of both eIF2α^S/S^ and eIF2α ^A/A^ cells (**Suppl Figure 1d**), showing that eIF2B’s ability to stimulate mutant KRAS signaling may be independent of its regulation by eIF2α phosphorylation at the level of translation.

These data highlight a distinct role for eIF2B in regulating MAPK signaling in cells expressing mutant versus wild-type KRAS and underscore its specific function in promoting MAPK activation and enhancing the survival of both human and mouse tumor cells harboring mutant KRAS.

### eIF2B binds to mutant, but not wild-type, KRAS 4B

We analyzed mass spectrometry (MS) data from experiments aimed at identifying KRAS-interacting proteins co-immunoprecipitated (co-IP) with FLAG-tagged wild-type KRAS and FLAG-tagged KRAS G12V in HEK293 cells (23). The analysis revealed that all five eIF2B subunits are in complex with the KRAS 4B isoform, but not KRAS 4A, only when KRAS 4B carries the G12V mutation (**Figure 2a**). We validated the interaction between endogenous mutant KRAS and eIF2B through co-IP experiments using extracts from H358 cells and antibodies targeting individual eIF2B subunits (**Suppl Figure 2a**).

**Figure 2.**
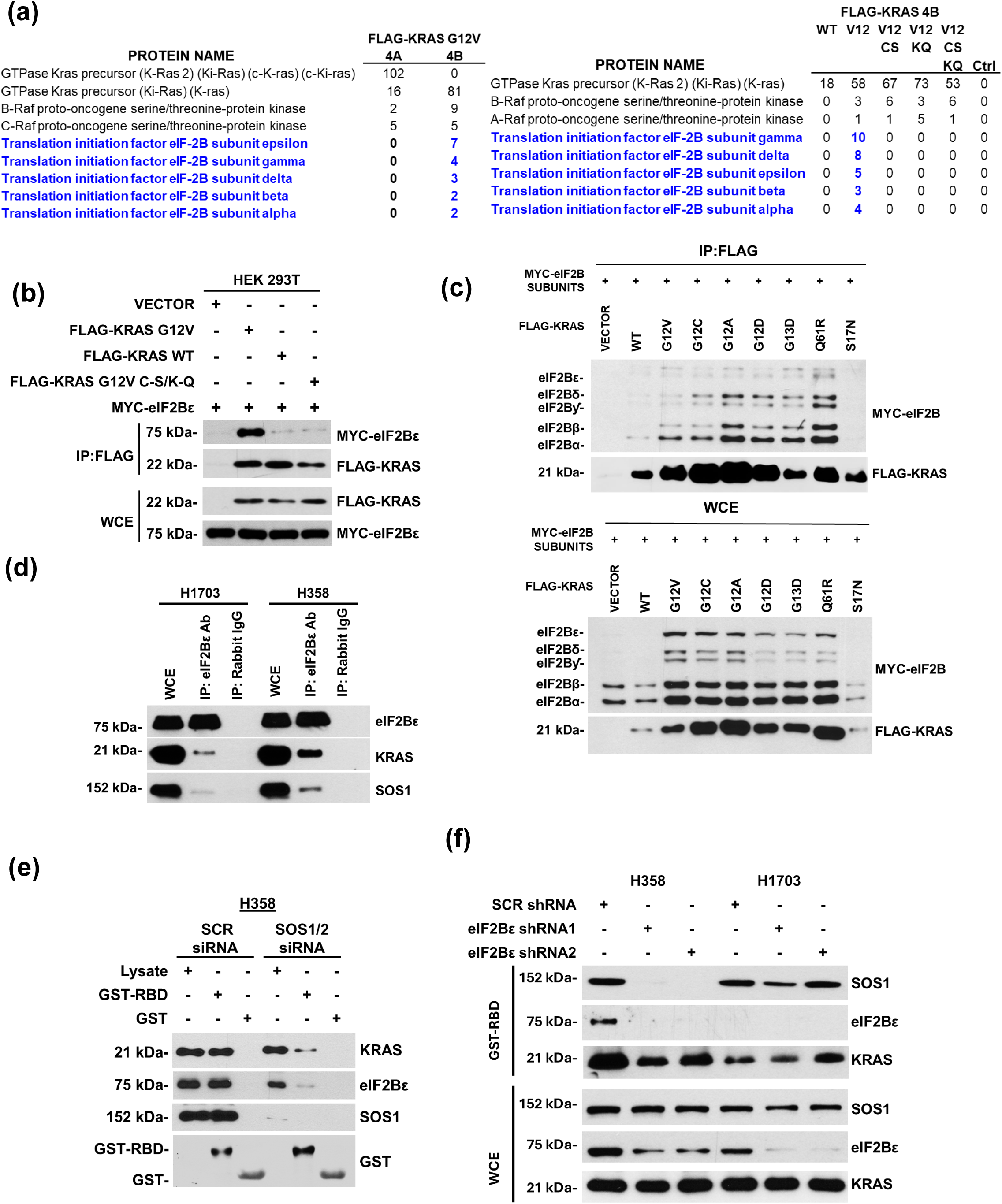
eIF2B physically interacts with GTP-bound mutant KRAS and SOS. (a) Mass spectrometry analysis of eIF2B and KRAS interactions. (Left panel) eIF2B subunits specifically interact with the G12V mutant of FLAG-KRAS 4B, but not with the G12V mutant of FLAG-KRAS 4A. (Right panel) Mutations in the C-terminus hypervariable region of FLAG-KRAS G12V impair its interaction with the eIF2B subunits. V12, G12V mutation of KRAS 4B; CS, C→S mutation in the C-terminal CAAX motif of KRAS G12V; KQ, K→Q mutations in the poly-lysine stretch of KRAS G12V; Ctrl, mass spectrometry analysis of proteins bound to a non-target antibody. (**b**, **c**) HEK293T cells were co-transfected with MYC-tagged constructs for eIF2Bε (panel b) or all eIF2B subunits (panel c), together with KRAS G12V variants containing C→S and polyK→polyQ mutations within the HVR (panel b) as well as various KRAS mutants harboring substitutions at G12, G13, or Q61 (panel c). Cell lysates were subjected to IP with an anti-FLAG antibody, followed by immunoblotting with anti-FLAG and anti-MYC antibodies to detect KRAS and eIF2B, respectively. Lysates from HEK293T cells transfected with insert less vector DNA served as negative controls. Protein loading in the co-IP assays was verified by immunoblotting of whole-cell extracts (WCE) (**d**) Interaction between endogenous eIF2B, SOS and KRAS in H358 and H1703 cells in co-IP assay with antibodies against the eIF2Bε subunit. Proteins were analyzed by immunoblotting to detect endogenous KRAS, SOS and eIF2Bε. Rabbit IgG was used as a negative control. Whole-cell extracts (WCE; 50 µg protein) were included as loading controls. (**e**, **f**) Cell lysates from H358 and H1703 cells were subjected to pull-down assays with GST-RBD of RAF. In panel (e), lysates from SOS1/2-proficient and SOS1/2 KD H358 cells were analyzed for bound KRAS, eIF2Bε, and SOS1 by immunoblotting. In panel (f), lysates from eIF2Bε-proficient and eIF2Bε-KD H358 (KRAS G12C) and H1703 (wild-type KRAS) cells were processed similarly using GST-RBD to detect KRAS, eIF2Bε, and SOS1. Whole-cell extracts (WCE; 50 µg protein) were included as loading controls.

MS analysis indicated that mutations in the carboxyl-terminal hypervariable region (HVR) of KRAS did not result in detectable interactions between FLAG-KRAS 4B G12V and endogenous eIF2B (**Figure 2a**). These mutations include the substitution of cysteine (C) for serine (S) in the CAAX motif or the polylysine (K)-to-polyglutamine (Q) substitution in HVR, which disrupt KRAS G12V binding to the PM (24). We observed a stronger interaction of FLAG-KRAS G12V and MYC-tagged eIF2Bε compared to FLAG-wild type KRAS in co-IP assays performed in HEK293 cells (**Figure 2b**). However, FLAG-KRAS G12V variants carrying a C-to-S mutation in the CAAX motif and poly K-to-poly Q mutations in HRV displayed a markedly reduced interaction relative to FLAG-KRAS G12V(**Figure 2b**).

Furthermore, co-IP assays of FLAG-KRAS 4B isoform in protein extracts from HEK293 cells revealed that KRAS G12V exhibited stronger interaction with endogenous eIF2B subunits compared to wild-type KRAS (**Suppl Figure 2b**). This interaction was notably reduced when KRAS G12V carried mutations in the HVR domain.

Moreover, incubation of protein extracts from HEK293 cells expressing MYC-tagged eIF2B subunits and FLAG-KRAS 4B with a biotinylated form of ISRIB, a small molecule that stabilizes the decameric structure of eIF2B (15), demonstrated the formation of an eIF2B:KRAS G12V complex in streptavidin-agarose pull-down assays (**Suppl Figure 2c**). This complex formation was not observed when FLAG-KRAS was wild type or G12V carried additional mutations in the HVR domain.

In additional experiments we tested the interaction between MYC-eIF2B subunits and different mutants of FLAG-KRAS 4B in HEK293 cells (**Figure 2c**). Notably, co-expression of FLAG-KRAS and MYC-eIF2B subunits led to increased levels of both KRAS and eIF2B in whole cell extracts, but only when mutant KRAS forms were co-expressed (**Figure 2c**). Given that the transfection experiments were normalized using *Renilla* luciferase as an internal control, these findings reveal that the co-expression of eIF2B and mutant KRAS may result in the mutual stabilization of these proteins. The co-IP assays revealed a strong interaction between MYC-eIF2B and FLAG-KRAS carrying mutations at G12 or Q61. In contrast, no interaction was detected with FLAG-KRAS WT or KRAS S17N, the latter of which has low affinity for GTP and is resistant to regulation by GEFs (25) (**Figure 2c**).

Collectively, the findings demonstrate that eIF2B specifically interacts with mature, PM-bound mutant KRAS and that enhanced GTP-binding of mutant KRAS is critical for this interaction.

### eIF2B recruits SOS in its interaction with mutant KRAS

Previous studies revealed structural parallels between eIF2Bε and SOS, the *bone fide* GEF of RAS proteins (26), particularly between helices III–VIII of eIF2Bε and residues 605–739 of SOS that form the regulatory RAS-GTP binding site (27), indicating a potential role for eIF2Bε in modulating SOS:RAS interactions We detected an interaction among endogenous SOS, eIF2Bε, and KRAS in co-IP assays. Notably, this interaction was stronger in H358 cells harboring KRAS G12C compared to H1703 cells expressing wild-type KRAS (**Figure 2d)**.

The interaction between SOS, eIF2B and KRAS G12V is direct, as demonstrated using purified recombinant proteins (**Suppl Figure 2d**). Specifically, we incubated the eIF2B β, γ, δ, and ε subunits with HIS-tagged KRAS G12V and a biotinylated FLAG-tagged SOS^cat^ encompassing residues 566-1046 of the regulatory (REM) and catalytic (CDC25) domains of SOS (28), followed by affinity purification using streptavidin-agarose beads. Immunoblot analysis confirmed the formation of an eIF2B(β,γ,δ,ε):KRAS G12V:SOS^cat^ complex, whose assembly was further enhanced by the addition of ISRIB, a small molecule known to stabilize eIF2B. These results support direct interaction among all three components in complex formation.

We next examined whether the interaction among eIF2B, SOS, and KRAS in cellular extracts depends on the presence of mature, lipid-bound KRAS. Co-IP experiments in HEK293 cells expressing MYC-eIF2B subunits revealed a robust interaction with HA-SOS in the presence of FLAG-KRAS 4B G12V (**Suppl Figure 2e**). In contrast, this interaction was markedly reduced when either wild-type KRAS or the KRAS G12V variant carrying mutations in the HVR were expressed, indicating that eIF2B:SOS association requires membrane-anchored KRAS.

We further examined whether the interaction of eIF2B with SOS also occurs with other RAS isoforms, including KRAS 4A, HRAS, and NRAS, in either their wild-type or G12V mutant forms. Co-IP assays in HEK293 cells revealed that KRAS 4B, particularly in its G12V form, exhibited a stronger interaction with eIF2Bε and SOS compared to HRAS, regardless of whether HRAS was wild-type or G12V (**Suppl Figure 2f**). When comparing the G12V forms of KRAS 4B, KRAS 4A, HRAS, and NRAS, KRAS 4B G12V displayed the strongest interaction with eIF2Bε and SOS (**Suppl Figure 2g**). These findings indicate that the eIF2B:SOS complex selectively recognizes structural or membrane-associated features unique to mutant KRAS 4B.

### eIF2B, via its interaction with SOS, stimulates the GTP-bound state of mutant KRAS

Mutant KRAS variants exhibiting intrinsic GTPase activity or low GDP dissociation rates retain partial dependence on SOS for GTP loading (29). The selective interaction of eIF2B with SOS and mutant KRAS led us to investigate whether eIF2B contributes to the formation of GTP-bound mutant KRAS.

To this end, we performed pull-down assays using a GST-fusion protein containing the RAS-binding domain of RAF (GST-RBD), a widely used tool for detecting GTP-bound, active RAS (30). In H358 cells harboring KRAS G12C, SOS1 and SOS2 KD impaired the association of mutant KRAS with GST-RBD and reduced the interaction of eIF2Bε with GST-RBD (**Figure 2e**). Notably, eIF2Bε KD via siRNA or shRNA reduced the binding of both KRAS and SOS to GST-RBD (**Figure 2f** and **Suppl Figure 2h**). In contrast, this effect was not observed in H1703 cells expressing wild-type KRAS (**Figure 2f**). These findings indicate that eIF2B specifically cooperates with SOS to promote the GTP-bound active conformation of mutant KRAS.

We further examined the effects of eIF2B stimulation by ISRIB on the GTP-bound state of KRAS in H358 cells. Our results demonstrated that the addition of ISRIB to protein extracts enhanced the binding of all five eIF2B subunits to GST-RBD (**Suppl Figure 2i**). Additionally, ISRIB promoted the interaction of SOS with GST-RBD, which was accompanied by an increased formation of the GTP-bound mutant KRAS (**Suppl Figure 2i**).

Conversely, treatment with BI-3406, a selective inhibitor of the SOS1:RAS interaction (31), decreased both GTP-bound KRAS levels and the recruitment of eIF2B, especially its catalytic subunit eIF2Bε, and SOS1 to GST-RBD of RAF in H358 cell extracts (**Suppl Figure 2j**). This inhibitory effect was reversed by ISRIB co-treatment, further supporting ISRIB’s role in stabilizing the eIF2B:SOS1:mKRAS complex. Following a similar approach, we observed that the addition of KRAS inhibitor, AMG510, a selective irreversible inhibitor of KRAS G12C (32), reduced the association of eIF2B subunits with GST-RBD, along with the inhibition of the GTP-bound state of KRAS (**Suppl Figure 2k**). However, these effects of AMG510 on eIF2B and mutant KRAS were reversed upon the addition of ISRIB.

Collectively, these findings demonstrate that eIF2B acts via SOS to promote the GTP-bound active state mutant KRAS in cells.

### Computational Modeling and Mapping of the eIF2B:SOS:KRAS Complex

To gain insight into the potential structural organization of the KRAS:SOS:eIF2B complex, we integrated existing structural data with AlphaFold 3 predictions (33). The resulting structural model (**Figure 3a** and **Suppl Figure 3)** provides two possible alternative configurations. In the first configuration, while KRAS occupies the catalytic site of SOS, the eIF2Bε subunit engages the allosteric RAS-binding site of SOS (**Figure 3a**), a site known to boost SOS’s GEF activity and promote GDP release from RAS at its catalytic site (28). In the second configuration, the eIF2Bε subunit interacts directly with a GTP-bound KRAS dimer (or higher-order oligomer), whereas another KRAS molecule binds the allosteric site of SOS (**Figure 3a**). Given the high symmetry of the eIF2B decamer, multiple interaction combinations are possible, including bivalent associations (eIF2B:SOS, eIF2B:RAS) and trivalent assemblies (RAS:eIF2B:SOS, RAS:eIF2B:RAS, SOS:eIF2B:SOS).

**Figure 3.**
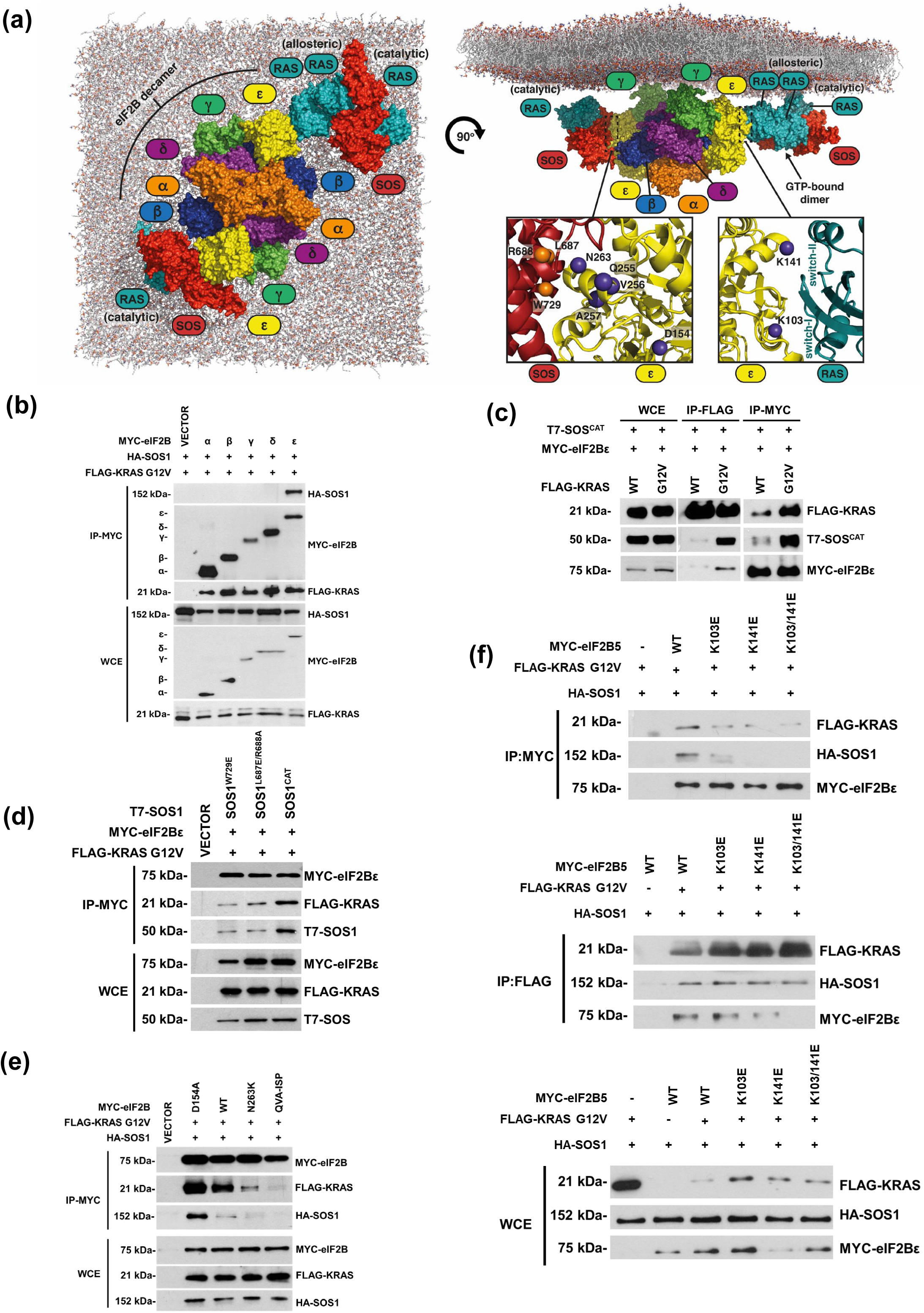
In silico assembly and biochemical mapping of the eIF2B:SOS:KRAS complex. (a) Putative model of the eIF2B:SOS:KRAS G12V structural assembly. eIF2B is predicted to associate either to the allosteric binding site of SOS or to GTP-bound RAS (dimer or oligomer) via its eIF2Bε subunit. In the close-up views of the predicted eIF2Bε interaction sites, the location of the mutated residues utilized in this study are highlighted with spheres on their Cα-atoms (eIF2Bε: blue spheres; SOS: orange spheres). For further information on the model see Suppl Figure 3. (b) The eIF2Bε subunit mediates the interaction of eIF2B with SOS and mutant KRAS. HEK293T cells were co-transfected with MYC-tagged constructs for each eIF2B subunit separately, HA-SOS1 and FLAG-KRAS G12V. Cell lysates were subjected to IP with an anti-MYC antibody, followed by immunoblotting with anti-FLAG, anti-HA, and anti-MYC antibodies to detect KRAS, SOS1, and eIF2B subunits, respectively. Protein loading in the co-IP assays was verified by immunoblotting of whole-cell extracts (WCE). (c) SOS residues 566–1046 (SOS^CAT^) interacts with eIF2Bε and mutant KRAS, but to a lesser extent wild type (WT) KRAS. HEK293T cells were transfected with MYC-tagged eIF2Bε, T7-tagged SOS^CAT^, and either FLAG-tagged KRAS G12V or WT KRAS. Cell lysates were co-IPed using anti-MYC or anti-FLAG antibodies, followed by immunoblotting with anti-MYC, anti-T7, and anti-FLAG antibodies. (d) Mutations in the allosteric RAS-binding site of SOS^CAT^ impair the interaction with eIF2Bε and mutant KRAS. HEK293T cells were transfected with MYC-tagged eIF2Bε, FLAG-tagged KRAS G12V and T7-tagged SOS^CAT^ either wild type or containing the W729E or L687E/R688A mutations. Cell lysates were co-IPed with anti-MYC antibodies, followed by immunoblotting with anti-MYC, anti-T7, and anti-FLAG antibodies to detect the respective proteins. (e) The catalytic GEF activity of eIF2Bε is essential for its interaction with SOS and mutant KRAS. HEK293T cells were transfected with HA-SOS1, FLAG-KRAS G12V and MYC-eIF2Bε either wild-type or carrying the hyperactive D154A mutation, the catalytically inactive N263K mutation, or the QVA→ISP mutation in the C-terminus. Cell lysates were co-IPed with anti-MYC antibodies, followed by immunoblotting with anti-MYC, anti-HA, and anti-FLAG antibodies to detect the respective proteins. (f) Mutations in eIF2Bε impair its interaction with SOS and mutant KRAS. HEK293T cells were transfected with FLAG-tagged KRAS G12V, HA-tagged SOS1, and MYC-tagged eIF2Bε, either wild-type or containing the K103E, K141E, or K103E/K141E mutations. Cell lysates were subjected to co-IP with anti-FLAG or anti-MYC antibodies, followed by immunoblotting with anti-MYC, anti-HA, and anti-FLAG antibodies to detect the respective proteins. (**b**-**f**) Data represent one of three reproducible experiments.

We tested the validity of this structural model by demonstrating that formation of the eIF2B:SOS:KRAS G12V complex is mediated by the catalytic eIF2Bε subunit, as shown by co-IP assays of individual MYC-eIF2B subunits with HA-SOS and mutant FLAG-KRAS in HEK293 cells (**Figure 3b**). Following a similar approach, we found that the eIF2Bε subunit exhibited a stronger interaction with T7-tagged SOS^cat^, which encompasses the regulatory REM and catalytic CDC25 domains of SOS (28), in the presence of FLAG-KRAS G12V compared with wild-type FLAG KRAS (**Figure 3c**). This interaction was mediated through the allosteric RAS-binding site of SOS, as the introduction of the W729E or L687E/R688A mutations (28) within this site markedly reduced complex formation (**Figure 3d**).

Moreover, substitution of Asp154 in MYC–eIF2Bε with a catalytic gain-of-function mutation (D154A), known to enhance GEF activity toward eIF2 (34), further strengthened its interaction with FLAG–KRAS G12V and HA–SOS in HEK293 cells (**Figure 3e**). In contrast, the catalytically inactive N263K mutation in MYC–eIF2Bε (34), as well as the ISP triple mutation (Q255I/V256S/A257P) within the C-terminal QVA motif, completely abolished association of MYC–eIF2Bε with both HA– SOS and FLAG–KRAS G12V (**Figure 3e**).

Notably, the SOS1 (W729E; L687E/R688A) and eIF2Bε (N263K; Q255I/V256S/A257P) mutations that disrupted complex formation map to residues located near the predicted SOS1:eIF2Bε interaction interface, whereas the D154A substitution in eIF2Bε, which did not impair complex assembly, is positioned distal to this interface (**Figure 3a**).

Introduction of eIF2Bε mutations K103E, K141E and K103E/K141E, predicted to be in eIF2Bε:KRAS interface (**Figure 3a**), reduced the formation of eIF2B:SOS:KRAS G12C complex in co-IP experiments in HEK293 cells (**Figure 3f**).

These results indicate that residues within the predicted eIF2Bε:KRAS and eIF2Bε:SOS1 interfaces are critical for the assembly of the eIF2B:SOS:KRAS complex, underscoring the structural specificity of eIF2Bε in mediating these protein–protein interactions.

### eIF2B co-localizes with SOS and mutant KRAS at PM

We performed immunofluorescence (IF) and confocal microscopy to investigate the subcellular localization of the eIF2B:SOS:KRAS complex in HEK293 cells co-expressing MYC-eIF2Bε, HA-SOS, and FLAG-KRAS G12V. Co-expression of all three proteins resulted in their accumulation at the cell periphery (**Figure 4a** and **Suppl Figure 4a**). This peripheral localization was enhanced by the expression of the hyperactive MYC-eIF2Bε D154A, which increased the number of cells displaying co-localization of the three proteins. In contrast, expression of the catalytically inactive MYC-eIF2Bε N263K markedly reduced this localization. Furthermore, introducing the ISP mutation, which disrupts the interaction between eIF2Bε and SOS (**Figure 3e**), also impaired peripheral localization of the complex (**Figure 4a**).

**Figure 4.**
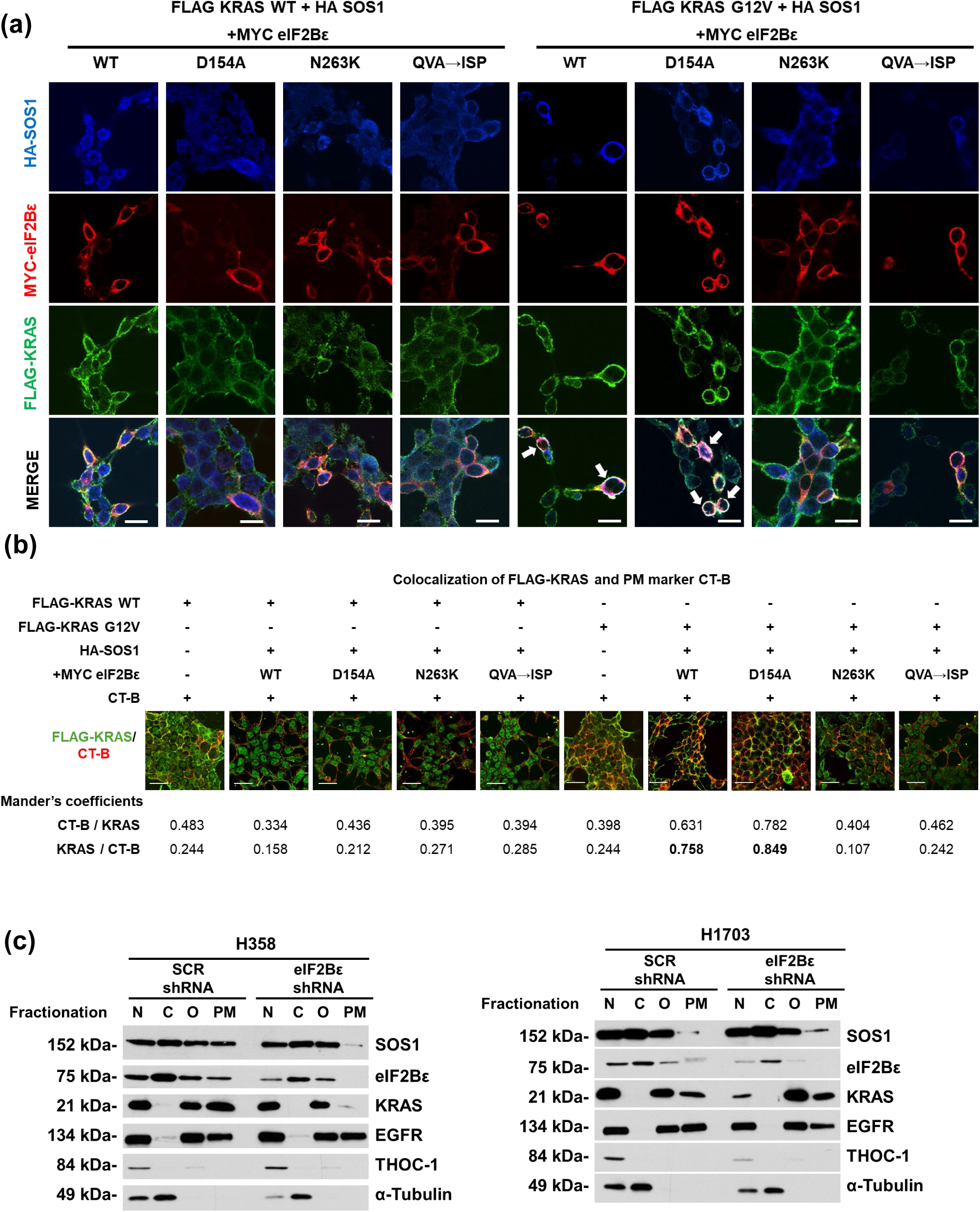
eIF2Bε co-localizes with SOS and mutant KRAS at the PM. (a) IF analysis of MYC-eIF2Bε (red, 568 nm), HA-SOS1 (blue) and FLAG-KRAS (green), either WT (left) or G12V (right), co-expressed in HEK293T cells. MYC-tagged eIF2Bε was expressed in HEK293T cells either as wild-type or in mutant forms, including the hyperactive D154A mutant, the catalytically inactive N263K mutant, or the QVA→ISP mutant that disrupts interaction with SOS. Protein localization was assessed by confocal microscopy. The white signal in the merged images indicates the co-localization of the three proteins at the periphery of the cell. (b) IF analysis of cells described in panel (a) assessing PM localization of FLAG-tagged KRAS (green) following co-staining with Cholera Toxin subunit B (CT-B, far-red, 647 nm), a PM marker. Mander’s colocalization coefficients, analyzed with JACoP (ImageJ), were used to quantify the extent of FLAG-KRAS colocalization with CT-B at the PM. Scale bar: 5 μm. The yellow signal in the images indicates the co-localization of FLAG-KRAS and CT-B at the PM. (c) H358 cells (KRAS G12C) and H1703 cells (wild-type KRAS) expressing either scrambled shRNA or eIF2Bε shRNA were fractionated into nuclear (N), cytosolic (C), organelle (O), and plasma membrane (PM) fractions. Each fraction was subjected to immunoblotting to detect SOS1, eIF2Bε, and KRAS. Fractionation quality was verified using specific markers: EGFR for the PM, THOC1 for the nucleus, and α-TUBULIN for the cytosol.

In contrast, substitution of KRAS G12V with wild-type KRAS markedly reduced the peripheral localization of MYC-eIF2Bε, HA-SOS, and FLAG-KRAS, highlighting the specificity of complex assembly and membrane targeting for the mutant KRAS (**Figure 4a** and **Suppl Figure 4b)**. Neither the hyperactive D154A nor the catalytically inactive N263K eIF2Bε mutants affected the localization pattern when wild-type KRAS was present. Similarly, the ISP mutation had no impact on protein localization in the context of wild-type KRAS (**Figure 4a**). Consistent with the localization data, expression of the hyperactive eIF2Bε D154A, but not the catalytically inactive eIF2Bε N263K, enhances ERK phosphorylation in HEK293 cells co-expressing KRAS G12V, but not in cells expressing wild-type KRAS (**Suppl Figure 4c**). These findings reveal that the ability of catalytically active eIF2Bε to promote PM localization of mutant KRAS is linked to its role in stimulating the MAPK pathway.

To evaluate PM localization of the eIF2B:SOS:KRAS complex, we stained cells with cholera toxin subunit B (CT-B), a well-established marker of the PM (35), and assessed its co-localization with FLAG-KRAS. Co-expression of HA-SOS with either wild-type MYC-eIF2Bε or the hyperactive MYC-eIF2Bε D154A significantly enhanced PM localization of FLAG-KRAS G12V (**Figure 4b**). In contrast, expression of the catalytically inactive MYC-eIF2Bε N263K mutant or the ISP mutant, which disrupts the interaction between eIF2Bε and SOS, markedly reduced FLAG-KRAS G12V localization at the PM (**Figure 4b**). In parallel experiments, expression of wild-type FLAG-KRAS resulted in substantially reduced PM localization, regardless of whether wild-type MYC-eIF2Bε or its mutants were co-expressed (**Figure 4b**).

When each MYC-tagged eIF2B subunit was expressed individually along with HA-SOS and FLAG-KRAS G12V, only the catalytic ε subunit was detected in the complex localized at the PM (**Suppl Figure 4d**). This observation aligns with co-IP data showing that eIF2Bε is essential for binding to both SOS and KRAS G12V (**Figure 3b**). Together, these findings highlight the requirement for the catalytic activity of eIF2Bε and its interaction with SOS in facilitating the PM localization of mutant KRAS.

To further substantiate these findings, we performed subcellular fractionation assays in KRAS G12C-expressing H358 cells and wild-type KRAS H1703 cells (**Figure 4c**). EGFR was used as a marker for the PM fraction, THO complex subunit 1 (THOC1) as a nuclear marker, and α-TUBULIN as a cytosolic marker. In H358 cells, both mutant KRAS and SOS were enriched in the PM fraction; however, this localization was markedly reduced upon KD of the eIF2Bε subunit, indicating a critical role for eIF2B in facilitating the membrane association of the KRAS:SOS complex. In contrast, eIF2Bε KD had no effect on the localization of wild-type KRAS and SOS in H1703 cells.

Interestingly, a portion of eIF2Bε, KRAS and SOS was also detected in the nuclear fraction of both cell lines. The presence of translation initiation factors in the nucleolus is likely related to their role in ribosome biogenesis (36), whereas the perinuclear localization of KRAS and nuclear localization of SOS has been reported earlier (37, 38). In H358 cells, eIF2Bε KD selectively reduced the PM-associated fraction of mutant KRAS without affecting its nuclear localization (**Figure 4c**). Conversely, in H1703 cells, eIF2Bε KD did not alter the PM-associated fraction of wild-type KRAS but led to a reduction in its nuclear fraction through an unknown mechanism (**Figure 4c**). Of note, we also observed a nuclear fraction of EGFR, consistent with studies reporting its nuclear presence in tumor cells (39).

Together, these data indicate that catalytically active eIF2B specifically promotes the PM localization of SOS and mutant KRAS, and that the interaction between eIF2Bε and SOS is essential for the assembly and PM targeting of the complex only in the presence of mutant KRAS.

### eIF2B facilitates SOS-mediated GDP-GTP exchange on RAS *in vitro*

Since the complex localizes at the PM, we next examined the mechanisms underlying the cooperative actions of eIF2B and SOS using an *in vitro* SOS-mediated RAS activation assay on supported membranes (40–43). This platform reconstitutes essential regulatory components— including full-length SOS, the membrane-linked scaffold linker for activation of T cells (LAT), the adaptor growth factor receptor-bound protein 2 (GRB2), phosphatidylinositol 4,5-bisphosphate (PIP2), and nucleotide-bound RAS to recapitulate the multistep regulation of SOS-mediated RAS activation (44). These steps include: (i) SOS recruitment to the membrane via GRB2; (ii) a PIP2-induced conformational change in SOS that exposes an allosteric RAS-binding site; (iii) release of SOS autoinhibition upon RAS binding; and (iv) SOS-catalyzed GDP–GTP exchange on RAS (**Suppl Figure 5a**).

Previous studies using this reconstituted system established that SOS autoinhibition release is a non-equilibrium stochastic process, with activation probability increasing as a function of membrane dwell time, a property governed by a protein condensation phase transition and kinetic proofreading mechanism (40, 42, 45).

Using this system, we systematically analyzed the recruitment dynamics of eIF2B to SOS:RAS-containing membranes to define its role in SOS-catalyzed RAS activation. To quantitatively assess membrane recruitment, we monitored the fluorescence intensity of mNeonGreen-labeled eIF2B decamers using total internal reflection fluorescence (TIRF) microscopy (**Suppl Figure 5b**). eIF2B exhibited slow, nonspecific recruitment to pLAT/GRB2-bound membranes (**Suppl Figure 5c**), and the addition of SOS did not alter this recruitment. In contrast, incorporation of RAS into the membrane markedly enhanced eIF2B recruitment by more than twofold compared with RAS-free membranes. Although SOS produced a modest additional increase, its effect was minor relative to that of RAS. Furthermore, eIF2B showed slightly stronger recruitment to GTP-bound RAS than to GDP-bound RAS, though this difference was limited. These results indicate that eIF2B binding to RAS is the principal determinant of its membrane association.

We examined whether eIF2B influences the catalytic rate of SOS-mediated RAS nucleotide exchange. Using a fluorescent RAS-GTP biosensor derived from the RAS-binding domain, we quantified the conversion of GDP-bound RAS to its GTP-bound form over time (**Suppl Figure 5d**). As expected, eIF2B alone did not catalyze nucleotide exchange, confirming that it lacks intrinsic GEF activity. SOS alone induced a concentration-dependent accumulation of RAS–GTP, consistent with previous reports (42, 46). Notably, the combination of SOS and eIF2B significantly increased the proportion of GTP-bound RAS compared with SOS alone, showing that eIF2B enhances SOS catalytic efficiency.

To further dissect this effect, we calculated the RAS nucleotide exchange rate per SOS molecule (**Suppl Figure 5e**), a measure that reflects the probability of SOS activation since the intrinsic catalytic rate of individual SOS molecules remains constant (47). Our analysis revealed that eIF2B substantially increased the exchange rate per SOS molecule as a function of the fraction of RAS in the GTP-bound state. These findings indicate that eIF2B promotes SOS autoinhibition release through GTP-bound RAS–dependent positive feedback, thereby accelerating RAS activation.

The *in vitro* data reveal that eIF2B binds RAS in a membrane-dependent fashion and amplifies SOS activity via GTP-bound RAS-mediated autoinhibition release. However, these findings raise the question of how eIF2B selectively activates mutant KRAS in cells, given that the *in vitro* SOS-mediated activation assay used HRAS, an isoform that is not affected by eIF2B in cellular contexts.

### eIF2B’s translational function connects glycosphingolipid biosynthesis to mutant KRAS

KEGG pathway analysis of the RNA-seq data from control and eIF2Bε KD H358 and MiaPaCa-2 cells (*KRAS G12C*) and H1703 cells (wild-type *KRAS*) revealed that the glycosphingolipid (GSL) biosynthesis pathway was upregulated by eIF2B in mutant KRAS cells but not in wild type KRAS cells (**Figure 5a**). Given that GSL production facilitates the proper localization of oncogenic KRAS to the PM (48), this pathway provides a mechanistic link through which eIF2B potentiates mutant KRAS activity at the PM.

**Figure 5.**
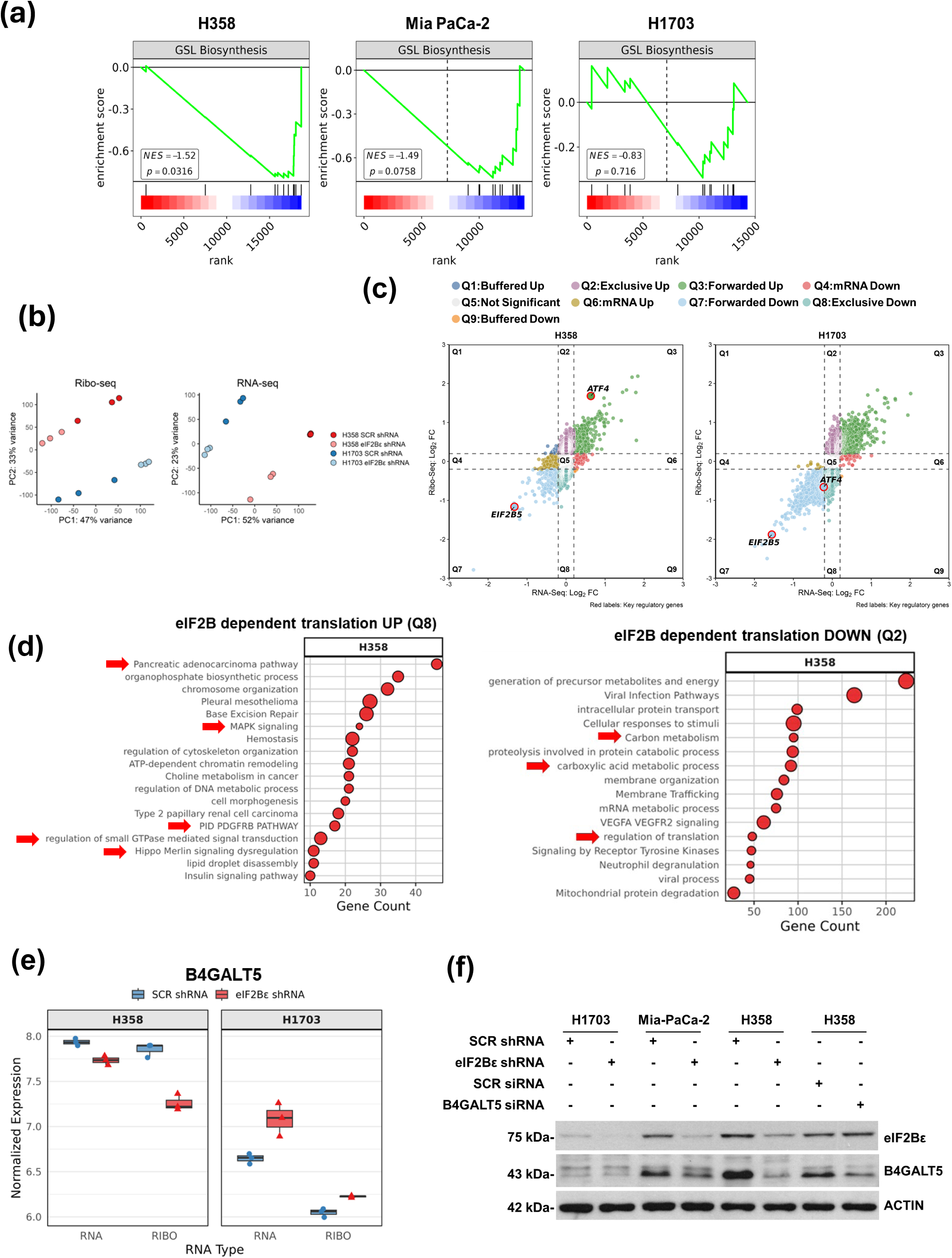
eIF2B promotes expression and translation of GSL biosynthesis genes. (a) KEGG pathway enrichment analysis of transcriptomic data from control and eIF2Bε KD cells. Pathway analysis revealed that eIF2Bε positively regulates GSL biosynthesis in H358 and MiaPaCa-2 cells containing *KRAS G12C*, but not in H1703 cells with wild-type *KRAS*. The negative enrichment scores for the GSL pathway following eIF2Bε KD in mutant *KRAS* cells indicate that eIF2Bε promotes transcriptional activation of genes involved in this signaling cascade (b) Principal-component analysis (PCA) plots of Ribo-seq (left) and RNA-seq (right) datasets separating each treatment and genotype. Each circle shows a biological replicate; different conditions and tumor types are color coded. (c) Scatterplot showing the association between RNA-seq and Ribo-seq fold change (FC) between H358 cells (left) and H1703 cells (right) treated with scrambled (SCR) shRNA and eIF2Be shRNA. mRNAs with statistically significant changes (p-adj < 0.05 and Log_2_ FC > 0.2 or Log_2_ FC <-0.2) are highlighted with different colors according to the following categories: Q1, buffered up, decrease in mRNA/increase in translation efficacy (TE); Q2, exclusive up, increase in ribosome-protected fragment (RPF) and TE, no change in mRNA; Q3, forwarded up, increase in mRNA and RPF at the same rate, no change in TE; Q4, mRNA down, no change in RPF and TE; Q6, mRNA up, no change in RPF and TE; Q7, forwarded down, decrease in mRNA and RPF at the same rate, no change in TE; Q8, exclusive down, decrease in RPF and TE, no change in mRNA; Q9, buffered down, increase in transcription/decrease in TE. (d) Bar graphs display statistically significant pathways identified from genes that are translationally upregulated or downregulated by eIF2B in H358 cells but not in H1703 cells. Only pathways that were significantly enriched in H358, but not in H1703, were retained for analysis. (e) Dot plot comparing the total mRNA (RNA) and ribosome-associated mRNA levels (RIBO) for *B4GALT5* mRNA in H358 and H1703 cells. Basal translation of *B4GALT5* is significantly elevated in mutant KRAS cells and is markedly reduced upon eIF2Bε silencing compared to wild type KRAS cells. (f) Extracts from eIF2Bε-proficient and eIF2Bε-KD cells were subjected to immunoblotting using antibodies against the indicated proteins. Protein extracts from H358 cells transfected with scrambled siRNA or B4GALT5-specific siRNAs were used as controls.

Although eIF2B likely influences GSL gene expression indirectly through its activation of KRAS signaling (48), our findings reveal an additional, direct effect of eIF2B on GSL production at the level of mRNA translation. Specifically, we assessed eIF2B’s impact on mRNA translation in H358 cells and H1703 cells by performing ribosome profiling (Ribo-seq) in combination with RNA-seq in both cell lines (49). Principal component analysis (PCA) and pairwise correlation of the Ribo-seq and RNA-seq datasets confirmed high reproducibility among biological replicates and clearly distinguished cells with intact versus impaired eIF2Bε function via shRNA-mediated KD (**Figure 5b**). We quantified eIF2B-dependent translational efficiency (TE) changes using the ΔTE (deltaTE) method in both cell lines (49–51). In H358 cells, loss of eIF2Bε increased the TE of 983 mRNAs (Q2 panel) and decreased the TE of 254 mRNAs (Q8 panel) (**Figure 5c** and **Suppl Table 1**). In contrast, in H1703 cells, 883 mRNAs exhibited increased TE (Q2 panel) and 1,428 mRNAs decreased TE (Q8 panel) following eIF2Bε KD (**Figure 5c** and **Suppl Table 1**).

GO analysis of genes in the Q2 (high TE upon eIF2B KD) and Q8 (low TE upon eIF2B KD) panels identified distinct biological pathways regulated by eIF2B in each cell line. Genes translationally stimulated by eIF2B in H358 but not in H1703 cells (Q8 panel) exhibit roles in MAPK signaling, small GTPase-mediated signal transduction, PDGFRB signaling, and HIPPO Merlin signaling dysregulation, processes known to be implicated in KRAS-driven tumorigenesis (52, 53) (**Figure 5d**). Genes translationally repressed by eIF2B in H358 but not in H1703 cells (Q2 panel) were associated with pathways involved in carboxylic acid metabolism, mRNA processing, and receptor tyrosine kinase signaling (**Figure 5d**).

Among the transcripts with TE specifically enhanced by eIF2B in H358 cells was *B4GALT5* mRNA (**Figure 5e**). While total *B4GALT5* mRNA levels remained unchanged, ribosome occupancy was reduced following eIF2Bε KD, indicating a defect in translation. *B4GALT5* encodes lactosylceramide synthase, a key enzyme in GSL biosynthesis to promote PM localization of mutant KRAS 4B and tumorigenesis in pancreatic cancer models (48). In line with these findings, immunoblot analysis showed increased B4GALT5 protein levels in eIF2Bε-proficient H358 and Mia-PaCa-2 cells harboring KRAS G12C, compared to eIF2Bε-KD cells. This regulatory effect of eIF2B was not observed in H1703 cells expressing wild-type KRAS (**Figure 5f**).

These findings show that eIF2B enhances the expression and translation of GSL biosynthesis genes, most notably the translation of B4GALT5, in a mutant KRAS-dependent manner.

### eIF2B controls mutant KRAS via GSL-dependent membrane organization

By enhancing GSL biosynthesis, which stabilizes and promotes nano-clustering of mutant KRAS at PM (48), eIF2B may reinforce mutant KRAS membrane localization and its downstream signaling. To evaluate this possibility, we examined eIF2B’s role in mutant KRAS distribution in T47D cells expressing GFP-KRAS G12V and stained with CellMask, a membrane-labeling dye. The extent of mutant KRAS redistribution to endomembrane compartments was quantified by calculating Manders’ coefficient, which represents the fraction of CellMask signal co-localizing with GFP–KRAS G12V (54). In eIF2B-proficient cells, GFP–KRAS G12V predominantly localized to the PM, whereas in eIF2Bε KD cells, it redistributed to intracellular compartments (**Figure 6a**). This shift was accompanied by significantly higher Manders’ coefficients in eIF2Bε-KD cells, indicating enhanced localization of GFP–KRAS G12V to the endomembrane fraction. In contrast, the PM localization of GFP–HRAS G12V in T47D cells remained unchanged upon eIF2Bε KD, demonstrating that the effect of eIF2B is specific to mutant KRAS and does not generalize to other RAS isoforms (**Figure 6a**).

**Figure 6.**
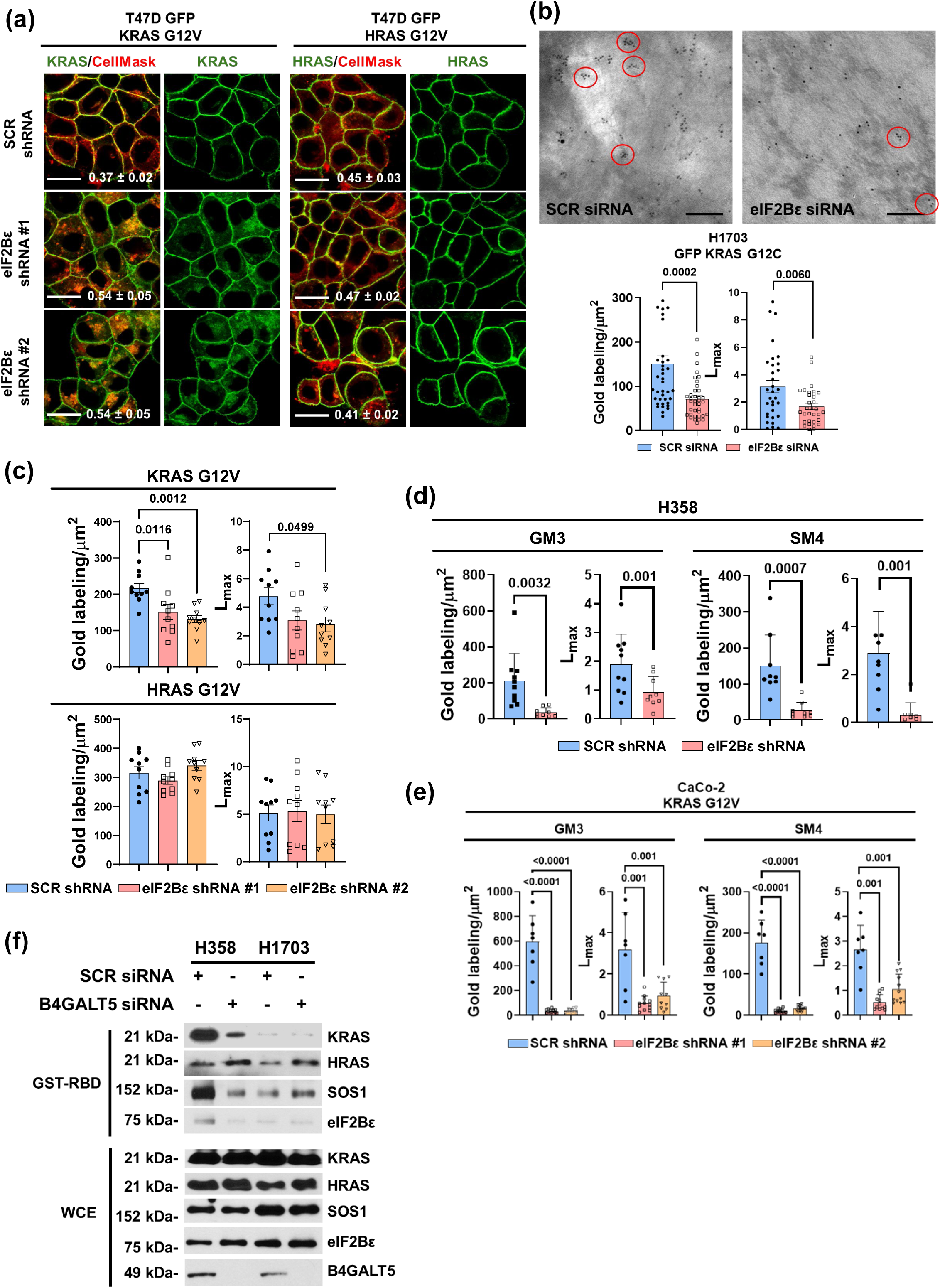
eIF2B supports mutant KRAS PM localization and nanoclustering via the GSL pathway. (a) eIF2B specifically promotes mutant KRAS localization at PM. Representative confocal images of T47D cells expressing either GFP-KRAS G12V or GFP-HRAS G12V, treated with either scrambled shRNA or eIF2Bε shRNA. Cells were stained with CellMask to label the PM. Co-localization of GFP-KRAS with CellMask was quantified using Manders’ coefficient and is presented as mean ± SEM (n = 3). Scale bar: 10 μm (b) eIF2B controls the localization and spatial organization of mutant KRAS at the PM. PM sheets were isolated from H1703 cells stably expressing GFP-KRAS G12C and transfected with either scrambled or eIF2Bε siRNA. The PM sheets were labeled with anti-GFP-conjugated gold particles and visualized by EM. Representative EM images are shown. Quantification of gold particles is presented as mean number ± SEM (n = 32). Spatial distribution was analyzed, and *L_max_* values, indicating the extent of KRAS G12C clustering, are shown in bar graphs (n = 32). Statistical significance was assessed using Student’s t-test for gold particle count (left) and bootstrap test for *L_max_* (right). Numeric values indicate *P*-values. Scale bar: 0.1 μm. (c) eIF2Bε depletion reduces mutant KRAS clustering. PM sheets were isolated from T47D cells stably expressing GFP-KRAS G12V or GFP-HRAS G12V along with eIF2Bε shRNA. The PM sheets were labeled with anti-GFP-conjugated gold particles and visualized via EM. The number of gold particles is presented as mean ± SEM (n = 10). Spatial mapping was also performed, and peak *L_max_* values, reflecting the degree of protein clustering, are shown as bar graphs. Numeric values indicate P-values. (**d**, **e**) eIF2Bε KD significantly reduces the PM levels of GM3 and SM4. PM sheets from H358 cells (KRAS G12C; panel d) or Caco-2 cells overexpressing GFP-KRAS G12V (panel e), treated with either scrambled shRNA or eIF2Bε-targeting shRNA, were fixed and labeled with 4.5 nm gold-conjugated anti-GM3 or anti-SM4 antibodies, then imaged by EM. Spatial distribution of gold particles was analyzed using univariate *K*-functions *(L(r) – r)*. PM levels of GM3 and SM4 were quantified as gold particle density per 1 μm², and clustering was assessed by the peak value of *L(r) – r* (*L_max_*). Statistical significance for labeling density and *L_max_* was determined using Student’s t-test and bootstrap analysis, respectively (n ≥ 12, mean ± SEM). (**f**) Silencing of B4GALT5 specifically reduces GTP-bound KRAS in mutant KRAS-expressing cells. H358 (KRAS G12C) and H1703 (WT KRAS) cells were transfected with either scrambled control or B4GALT5 siRNA. Protein extracts were subjected to pull-down assays using GST–RBD of RAF, followed by immunoblotting with antibodies against KRAS, HRAS, eIF2Bε, SOS1, and B4GALT5. Protein loading was assessed by immunoblotting of whole-cell extracts (WCE).

To directly quantify the dissociation of mutant KRAS from the PM, we prepared intact basal PM sheets from eIF2Bε-proficient and eIF2Bε KD H1703 cells expressing GFP–KRAS G12C. These membrane sheets were labeled with gold-conjugated anti-GFP antibodies and analyzed via electron microscopy (EM) (**Figure 6b**). Spatial distribution analysis of GFP–KRAS G12C using univariate K-function analysis revealed a marked decrease in *L_max_*, the peak of the *L(r)–r* clustering function, indicative of impaired nanoclustering (**Figure 6b**).

To assess whether the regulatory role of eIF2B extends beyond mutant KRAS to other RAS isoforms, we conducted similar EM analyses on T47D cells expressing GFP–KRAS G12V or GFP– HRAS G12V, with or without eIF2Bε KD. In GFP–KRAS G12V–expressing cells, eIF2Bε KD led to a significant reduction in PM-associated gold labeling and a substantial decrease in *L_max_*, indicating disrupted localization and nanoclustering (**Figure 6c**). In contrast, GFP–HRAS G12V localization and clustering were unaffected by eIF2Bε KD, as reflected by unchanged gold particle density and *L_max_* values (**Figure 6c**). Collectively, these findings demonstrate that eIF2B is specifically required for the proper PM localization and nanoclustering of mutant KRAS, a process essential for effective RAS signaling, and does not exert similar effects on other RAS isoforms such as HRAS.

The spatial organization of mutant KRAS at the PM is shaped by the synthesis of specific GSLs, particularly ganglioside GM3 and sulfatide SM4, in the outer leaflet (48). Given that our RNA-seq and Ribo-seq analyses revealed eIF2B-dependent upregulation of the GSL biosynthesis pathway (**Figure 5**), we investigated whether eIF2B controls the production of GM3 and SM4. Silencing eIF2Bε markedly reduced GM3 and SM4 levels at the PM in H358 cells as well as in Caco-2 cells overexpressing GFP-KRAS G12V (**Figure 6d,e**). These findings demonstrate that eIF2B promotes mutant KRAS plasma-membrane localization and nanoclustering by driving GSL biosynthesis.

We also assessed the influence of GSL pathway on the interaction of eIF2B with SOS and mutant KRAS. Pull-down assays using GST–RBD of RAF revealed that silencing of *B4GALT5* disrupted the formation of the eIF2Bε:SOS:KRAS complex in H358 cells (KRAS G12C), accompanied by a marked reduction in GTP-bound mutant KRAS levels (**Figure 6f**). Notably, GTP-bound HRAS levels were unaffected by B4GALT5 KD in H358 cells, and no comparable effects were observed in H1703 cells expressing wild-type KRAS. These findings demonstrate that eIF2B-driven GSL synthesis is required for the assembly of the eIF2B:SOS:KRAS complex and for maintaining GTP loading of mutant KRAS.

### eIF2B is a tumor promoter and prognostic marker in mutant KRAS cancer

The transforming activity of mutant KRAS requires SOS activity for full biological function (29). Given eIF2B’s role in stimulating SOS GEF activity toward mutant KRAS, we assessed its role in mutant KRAS-driven tumorigenesis in xenograft assays in immune deficient mice using H358 cells (*KRAS G12C*) and H1703 cells (wild-type *KRAS*). While eIF2Bε KD significantly suppressed the growth of H358 tumors (**Figure 7a**), it had no appreciable effect on H1703 tumor growth (**Figure 7b**), supporting a mutant KRAS-specific function of eIF2B in promoting tumor growth.

**Figure 7.**
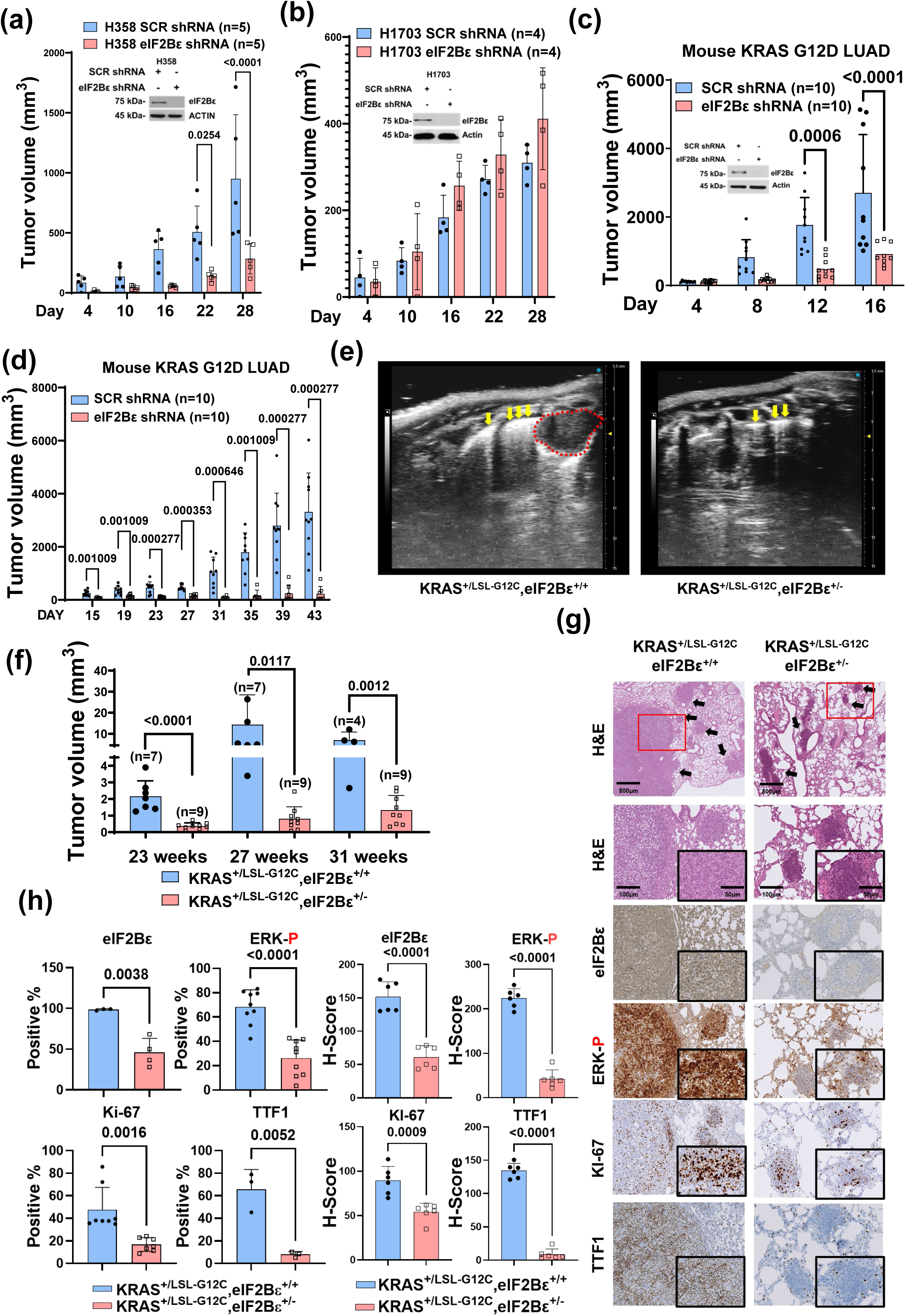
eIF2B promotes the growth of mutant KRAS-driven cancers. (**a**, **b**) H358 cells harboring KRAS G12C (panel a) and H1703 harboring wild type KRAS (panel b) were transduced with scrambled shRNA (control) or eIF2Bε shRNA and subcutaneously injected into immunodeficient nu/nu mice (H358 cells, n = 5; H1703 cells, n=4). Tumor size (mm³) was monitored over time. (**c**, **d**) Mouse KRAS G12D LUAD cells expressing either a scrambled shRNA or eIF2Bε shRNA were subcutaneously transplanted into immunodeficient nu/nu mice (panel c, n=10) and immunocompetent syngeneic C57BL/6 mice (panel d, n=10). (**e**, **f)** Mice expressing KRAS G12C and lacking TP53 in the lungs, with either intact eIF2Bε or heterozygous deletion eIF2Bε^+/−^, were monitored for tumor formation using ultrasound imaging to detect lung tumors located peripherally in the septum and in contact with the pleura. Representative ultrasound images of lung tumors at 27 weeks of tumor development are shown in panel (e), with tumor location indicated by arrows and tumor size marked by yellow dashed lines. Scale bars, 100 μm. Quantification of tumor growth over time based on ultrasound imaging is presented in panel (f). (**g**, **h**) IHC analysis of mouse lung tissue. Hematoxylin and eosin (H&E) staining and immunohistochemical (IHC) staining of lung tumors for eIF2Bε, phosphorylated ERK, Ki-67, and TTF1 were performed at 31 weeks following CRE-lentivirus intubation (panel g; n = 2 mice per genotype). Graphs represent the average H-score per tumor per lung section from mice expressing KRAS G12C with either eIF2Bε^+/+^ or eIF2Bε^+/−^ genotypes. Scale bars in H&E stained core tumor images correspond to 800 μm and 100 μm, respectively, and 50 μm in magnified images. (**a**-**d**, **f**, **h**) Quantification is presented as mean ± SD; P-values from Student’s t-tests are shown for significant differences only.

We further examined the tumor-promoting role of eIF2B using xenograft transplantation assays in mice, employing LUAD cells derived from a KRAS G12D-expressing, TP53-deficient mouse model (21). These KRAS G12D LUAD cells were engineered to express either a control non-targeting shRNA (PLKO) or an shRNA targeting eIF2Bε and subsequently transplanted into both immunodeficient mice and immune-competent syngeneic C57BL/6 mice. Our results demonstrated that eIF2Bε KD significantly impaired KRAS G12D-driven tumor growth in both models (**Figure 7c,d**). Notably, the inhibitory effect was more pronounced in immunodeficient mice (**Figure 7d**), indicating a potential interaction between eIF2B function in the tumors and regulation of immune responses in the tumor microenvironment (TME). These findings highlight a cell-intrinsic role for eIF2B in supporting mutant KRAS tumor growth and raise the possibility that eIF2B may modulate TME-associated pathways to enhance its tumorigenic potential.

To further support these findings, we established a genetically engineered mouse model of LUAD to assess the tumorigenic contribution of eIF2B *in vivo*. Mice harboring a *loxP-STOP-loxP* (LSL)-*KRAS G12C* allele were crossed with conditional eIF2Bε^f/f^ mice. Lung tumors were initiated via intratracheal instillation of lentiviruses co-expressing CRE recombinase and an shRNA targeting TP53 (21).

Ultrasound imaging of infected mice revealed that heterozygous deletion of eIF2Bε^+/−^ significantly reduced both the presence and size of lung lesions induced by KRAS G12C expression and TP53 inactivation at different time points post-intubation, compared to mice with eIF2Bε^+/+^ tumors (**Figure 7e,f**).

To better assess tumor growth, we performed immunohistochemical (IHC) analysis on lung tissue sections collected after 23 weeks post-intubation, a time point at which adenocarcinomas predominate in KP models (21). Immunostaining for hematoxylin and eosin (H&E) and marker of proliferation Kiel 67 (KI-67) indicated that mice with intact eIF2Bε^+/+^ developed distinct lung adenocarcinomas in response to KRAS G12C expression and TP53 loss, tumors were barely detectable in eIF2Bε^+/-^ mice (**Figure 7g,h**). KI-67+ lesions in eIF2Bε^+/+^ lungs also exhibited higher levels of eIF2Bε and phosphorylated ERK compared to eIF2Bε^+/−^ sections (**Figure 7g,h**). In addition, immunostaining for thyroid transcription factor 1 (TTF1), a marker of alveolar epithelial cells (55), revealed stronger expression in eIF2Bε^+/+^ than in eIF2Bε^+/−^ lung tissues (**Figure 7g,h**). Together, these findings highlight a critical role for eIF2B in promoting tumorigenesis in mutant KRAS-driven LUAD.

To assess the clinical relevance of our findings, analysis of mRNA expression data from The Cancer Genome Atlas (TCGA) revealed that elevated expression of the catalytic eIF2Bε subunit correlates with poorer prognosis in LUAD tumors harboring mutant, but not wild-type, KRAS (**Suppl Figure 6a**). This trend was further confirmed through examination of TCGA pan-cancer datasets, where the negative prognostic impact of high eIF2Bε expression was specifically observed in tumors with mutant *KRAS* (**Suppl Figure 6b**). The remaining eIF2B subunits do not exhibit the same prognostic significance as eIF2Bε (**Suppl Figure 7, 8**), except for eIF2Bβ, for which high mRNA expression is associated with reduced survival probability in TCGA pan-cancer datasets from tumors harboring mutant KRAS (**Suppl Figure 8**). These data highlight the potential of the catalytic eIF2Bε subunit as a clinically relevant marker for risk stratification in mutant KRAS cancers.

## DISCUSSION

### Non-Canonical Role of eIF2B in Mutant KRAS Activation

Our study uncovers a non-canonical role of eIF2B in activating mutant KRAS via its complex formation with SOS at PM, thereby enhancing KRAS signaling and tumorigenesis. Conceptually analogous, but mechanistically distinct from eIF2B, are the non-canonical activities of the GAP proteins RGS3 and RAN-GAP in regulating KRAS signaling (4, 37), highlighting how unconventional regulatory mechanisms can profoundly influence KRAS oncogenic function and reveal potential therapeutic opportunities.

### Assembly of the eIF2B:SOS:KRAS Complex

Biochemical data and computational modeling of the eIF2B:SOS:KRAS complex, informed by structural data from selected eIF2B and RAS complexes, supports the interaction between eIF2B and SOS through the allosteric binding site of SOS. The structural modeling indicates two possible KRAS:SOS:eIF2B arrangements: First, KRAS bound to SOS’s catalytic site with eIF2Bε occupying the SOS allosteric site, and second, eIF2Bε subunit interacting directly with a GTP-bound KRAS dimer or oligomer that is associated with the SOS allosteric site. Mutation mapping supports these interaction interfaces, and the symmetric architecture of the eIF2B decamer permits multiple (bi-or tri-valent) binding combinations. However, resolution of the model limits more detailed predictions or preferences among these protein-protein interactions. This structural versatility may contribute to KRAS nanoclustering at the PM involving the heterodimerization of mutant and wild-type KRAS. Such multivalent interactions and nanoclustering could also explain the observed co-precipitation of the eIF2B:SOS:KRAS complex in pull-down assays of GST-RBD RAF, given that the KRAS switch regions cannot simultaneously engage SOS, eIF2B, and RBD of RAF.

### Mechanistic Insights into SOS Activation by eIF2B

How does eIF2B, through its interaction with SOS, facilitate the engagement or stabilization of GTP-bound mutant KRAS within the eIF2B:SOS:RAS complex? KRAS mutants exhibiting intrinsic GTPase activity or low GDP release rates are only partially loaded with GTP and thus retain partial dependence on SOS for nucleotide exchange, though to a lesser extent than wild-type KRAS (29). One possibility, as instructed by computational modeling, is that eIF2B recruits and stabilizes SOS in active conformation. eIF2B, particularly through its catalytic ε subunit, binds to SOS within the allosteric RAS-binding domain, the same region normally engaged by GTP-bound RAS. This binding likely mimics or stabilizes the allosteric activation state of SOS. Consequently, eIF2B enhances the GEF activity of SOS, facilitating GDP release from mutant KRAS. Once stabilized by eIF2B in its active conformation, SOS can more efficiently catalyze GDP–GTP exchange on nearby mutant KRAS molecules as well as on wild-type KRAS molecules heterodimerized with mutant KRAS (56). For mutant KRAS variants that are constitutively GTP-bound, such as KRAS with Q61 mutations, eIF2B-mediated stimulation of SOS activity and its interaction with mutant KRAS may impact the activation of wild-type KRAS through heterodimerization and formation of KRAS nanoclusters. Through its interaction with eIF2B, SOS exhibits enhanced stability and GEF activity toward mutant KRAS, leading to increased accumulation of GTP-bound active KRAS.

### Specificity of eIF2B Toward Mutant KRAS 4B

The SOS-mediated RAS activation assay using recombinant proteins provides proof of concept that eIF2B functions as a GEF co-factor of SOS for RAS. In contrast, our experiments using cellular proteins reveal that the eIF2B:SOS:RAS complex forms far more robustly with KRAS 4B than with KRAS 4A, HRAS, or NRAS, in both wild-type and G12V mutant forms. Although the G-domains of Ras proteins share high structural similarity, intrinsic isoform-specific features, such as differences in helix α5 (57), may contribute to eIF2B’s preferential engagement with KRAS 4B, particularly in its mutant state.

Post-translational differences among RAS isoforms further refine this specificity. eIF2B selectively targets mutant KRAS 4B, but not the alternatively spliced KRAS 4A isoform. Although both isoforms share comparable oncogenic potential, they can operate through distinct mechanisms, for example, by differentially stimulating aerobic glycolysis via the induction and activation of key glycolytic enzymes, including hexokinases (58, 59).

A central determinant of this specificity is the unique mode of PM attachment used by KRAS 4B. Whereas KRAS 4A, HRAS, and NRAS depend on palmitoylation-mediated membrane anchoring via similar HVRs, KRAS 4B relies on a polybasic domain (24). We find this domain to be essential for the interaction of mutant KRAS 4B with eIF2B and SOS. PM lipid composition is known to regulate RAS nanoclustering through interactions between C-terminal membrane anchors and specific phospholipids (60). Consistent with this principle, we show that eIF2B binds mature, membrane-anchored mutant KRAS 4B and promotes its nanoclustering at the PM. These findings indicate that PM lipid composition is a key factor underlying eIF2B’s specificity for mutant KRAS 4B over other RAS proteins.

What accounts for eIF2B’s preferential engagement with mutant over wild-type KRAS 4B? Our data indicate that this selectivity stems from eIF2B’s ability to enhance GSL synthesis, particularly GM3 and SM4, in the outer leaflet of the PM. Increased levels of these GSLs promote the redistribution of phosphatidylserine within the inner leaflet, generating a membrane environment that preferentially supports the anchoring and function of mutant KRAS 4B (48, 61). Mechanistically, eIF2B drives this process by elevating the expression of GSL biosynthetic genes through the stimulation of mutant KRAS signaling and by translationally upregulating *B4GALT5* mRNA, which encodes a key GSL-producing enzyme, selectively in mutant KRAS-expressing cells. Silencing B4GALT5 disrupts the eIF2B:SOS:KRAS complex and reduces GTP-loaded mutant KRAS, without affecting HRAS or wild-type KRAS.

Thus, eIF2B-dependent lipid remodeling via the GSL pathway supports nanoclustering and membrane localization of mutant KRAS 4B, defining a mechanism by which eIF2B selectively activates mutant KRAS.

### Subcellular Localization and eIF2B Bodies

We demonstrate that eIF2Bε co-localizes with SOS and mutant KRAS 4B at the PM, but not with wild-type KRAS 4B. In yeast and mammalian cells, eIF2B forms cytoplasmic foci called eIF2B bodies that serve as sites of GEF activity for eIF2 (62, 63). The formation and organization of these bodies are regulated in a cell type-and stress-specific manner (62, 64). Given the dynamic subcellular distribution of eIF2B under different conditions, its localization to the PM may involve coordinated actions among its subunits to support the anchoring and activation of mutant KRAS at the membrane. Further studies are needed to clarify the role of eIF2B body formation in regulating SOS and mutant KRAS at the PM, as well as how this process is influenced by cellular stress.

### Translational Regulation and Oncogenic Signaling

A defining feature of eIF2B is its ability to regulate mRNA translation initiation through its GEF activity on eIF2 (15). During stress, eIF2B is essential in translational regulation of adaptive mRNAs via eIF2 phosphorylation dependent as well as independent pathways (15, 20). Our investigation of eIF2B’s translational function in non-stressed cells reveals its role in promoting oncogenic pathways downstream of mutant KRAS. Notably, the enhanced translation of *B4GALT5* mRNA represents a key mechanism by which eIF2B upregulates GSL synthesis, thereby facilitating the anchoring and activation of mutant KRAS at PM.

Our data from mouse LUAD cells that were either proficient or deficient in eIF2α phosphorylation show that eIF2B stimulates mutant KRAS signaling independently of phosphorylated eIF2. These findings reveal that the translational effects of phosphorylated eIF2, mediated through its inhibition of eIF2B, are mechanistically distinct from eIF2B’s role in stimulating mutant KRAS activity. However, it remains to be determined whether cellular stress alters the balance between eIF2B in complex with SOS and mutant KRAS at the PM versus eIF2B bound to eIF2 in tumors, thereby modulating mRNA translation.

### Functional and Clinical Implications

Mutant KRAS requires SOS activity for full biological function in tumor cells (29). The ability of eIF2B to stimulate mutant KRAS activity via SOS and downstream effector MAPK and GSL biosynthesis pathways underlies its tumorigenic potential in cancer models.

We show that eIF2B promotes tumorigenesis in human tumor cells expressing mutant KRAS, but not wild-type KRAS, in xenograft assays. Furthermore, analysis of an autochthonous mouse model of KRAS G12C-driven LUAD highlights the essential role of eIF2B in promoting lung tumorigenesis. Clinically, high expression of the eIF2Bε is associated with poorer outcomes in patients bearing KRAS-mutant tumors, supporting its contribution to tumor progression and its potential prognostic value. These results underscore the critical contribution of eIF2B to mutant KRAS-driven tumorigenesis and its potential as a therapeutic target.

## Conclusion

Our study identifies eIF2B as a novel regulator of mutant KRAS activation and tumorigenesis. By forming a complex with SOS and mutant KRAS at the PM, eIF2B enhances SOS catalytic activity, promotes KRAS GTP loading, and drives oncogenic signaling. Through its additional role in regulating GSL synthesis and translation, eIF2B facilitates KRAS 4B membrane clustering and activation, linking translational control to membrane signaling. These findings establish eIF2B as a key modulator of mutant KRAS-driven cancers and a potential therapeutic target.

## ACKNOWLEDGEMENTS

We acknowledge the expertise of Naciba Benlimame and Marie-Lyne Fillion at the core facility of the *Lady Davis* Institute for assistance in immunohistochemistry and cell imaging services. We thank Christopher Proud for MYC-eIF2B constructs; Dafna Bar-Sagi for SOC^Cat^ constructs; Dominique Esposito for NRAS, HRAS and KRAS constructs; Jothi Krishnamoorthy for technical support. This work was funded by grants from the Canadian Institutes of Health Research (CIHR; PJT-168864) and Canadian Cancer Society Research Institute (CCSRI) Innovation grant awarded to AEK. HK and SD are recipients of the Faculty of Medicine awards from McGill University, and JYZ is supported by a studentship award from *Le Fonds de recherche du Québec* (*FRQS*). We thank CSC– IT Center for Science, Finland, for computational resources.

## AUTHOR CONTRIBUTIONS

Conceptualization: AEK, HK, K-JC, JL, MH, JTG, NS, PW. Methodology: HK, K-JC, H-RL, JL, PE, NG, SW, JYZ, MA, RG, KMR, QD, AS, JA, BT, RL, CRA-N, J-PL, JC, CBM. Software: SD, TP, MK, JA. Validation: K-JC, NG, RG,KMR, BT, JC. Formal Analysis: HK, SD, K-JC, H-RL, JL, RL, JC, TJB, CBM, MI, JTG, NS, PW, AEK. Investigation: HK, SD, K-JC, H-RL, JL, PE, TP, MK, NG, MA, AS, BT, RL, JC, CBM. Data Curation: HK, SD, K-JC, H-RL, JL, TP, MK, MA, AEK. Writing: HK, AEK, K-JC, H-RL, JL, PE, TP, MK, J-PL, AEK. Visualization: HK, SD, K-JC, H-RL, JL, TP, MK. Supervision: AEK, K-JC, JL, TP, JH, DL, TJT, MI, JTG, NS, PW. Project Administration: AEK.

## DATA AVAILABILITY STATEMENT

The RNA seq data generated in this study have been deposited in the Gene Expression Omnibus (GEO) GSE287513. Structures of the generated models are provided in SI files. AlphaFold 3 predictions: Non-commercial use only, subject to AlphaFold Server Output Terms of Use (https://alphafoldserver.com/output-terms); no use in docking or screening tools.

## METHODS

### Cell lines

H358, NCI-H1703, H2122 and T47D cells were maintained in Roswell Park Memorial Institute-1640 (RPMI-1640, Wisent) supplemented with 10% fetal bovine serum (FBS; Wisent) and 100 U/mL penicillin–streptomycin (Wisent). T47D cultures received an additional 10 µg/mL human recombinant insulin (Gibco, Cat#12585-014). HCT116, HK2-8, MiaPaCa-2, and HEK293T cells were cultured in Dulbecco’s Modified Eagle Medium (DMEM; Wisent) with 10% FBS and 100 U/mL penicillin–streptomycin. Caco-2 cells were grown in Minimum Essential Medium (MEM; Wisent) containing 20% FBS. Primary KRAS G12D eIF2α ^S/S^ and eIF2α ^A/A^ lung tumor cells were established as described (21) and maintained in RPMI-1640 supplemented with 10% FBS, 100 U/mL penicillin– streptomycin, 0.075% NaHCO₃ (Life Technologies), 1× essential amino acids, and 1× non-essential amino acids (Life Technologies). All cultures were incubated at 37 °C in a humidified 5% CO₂ atmosphere. The cells were free of mycoplasma contamination.

### Stable eIF2Bε knockdown

H358, H1703, MiaPaCa-2, H2122, and T47D cells expressing GFP–KRAS G12V or GFP-HRAS G12V, as well as Caco-2 cells expressing GFP–KRAS G12V, were transduced with pLKO.1 lentiviral particles encoding either a non-targeting (scrambled) shRNA or an shRNA specific for human eIF2Bε. Forty-eight hours post-infection, cells were selected in growth medium containing 2.5 µg/mL puromycin until all non infected cells were eliminated. Stable knockdown pools were maintained in 2.5 µg/mL puromycin for all downstream experiments. Colony formation assays of eIF2Bε Knockdown cells were conducted with 2×10^3^ cells in 6-well plate for up to 10 days. Cells were fixed in 3.7% formaldehyde (v/v) and stained with 0.2% crystal violet (w/v). Colonies were scored using an automated cell colony counter (GelCount; Oxford Optronix), analysed with ImageJ 1.54g software (NIH, Maryland, USA).

### Transient gene knockdown

Transient downregulation of eIF2Bε, SOS1/2 and B4GALT5 was achieved using four different siRNAs (Horizon Discovery) with sequences listed in Supplementary Table 1. 2.5 × 10^5^ cells in 60-mm tissue culture plates were transfected with 200 pmol siRNA for target protein or scrambled sequence siRNAs (Dharmacon) using Lipofectamine 2000 (Invitrogen) following the manufacturer’s instructions. Cells were refreshed with appropriate medium supplemented with 10% FBS and 1% antibiotics after 4 hours of incubation and incubated at 37 °C for 48 hours before further analysis.

### DNA transfection

DNA transfections were carried out using Lipofectamine 2000 reagent (Invitrogen) according to the manufacturer’s specifications. Cells were refreshed with appropriate medium supplemented with 10% FBS and 1% antibiotics after 4 hours of incubation and incubated at 37 °C for 48 hours before further analysis. Plasmids encoding Myc–His–tagged human eIF2B subunits (α, β, γ, δ, ε) and FLAG– His–tagged KRAS WT, G12V, G12V/C-S, and G12V/C-S/K-Q constructs were described previously (23). HA-SOS1 were sourced from AddGene (Plasmid #32920). Site-directed mutagenesis was performed (Mutagenex) to generate the following variants: eIF2Bε point mutants (D154A, N263K, and Q^255^V^256^A^257^→I^255^S^256^P^257^) and FLAG–His–tagged KRAS mutants (G12C, G12A, G12D, G13D, Q61R, S17N). Truncated SOS variants (SOS^cat^, SOS^W729E^, and SOS^L687E/R688A^) were kindly provided by Dr. Dafna Bar-Sagi (New York University, School of Medicine). *Firefly* luciferase assays were performed with the Dual-Luciferase Reporter Assay System (Promega) using *Renilla* luciferase reporter gene serving as an internal control (65).

### Immunoblotting

Cells were washed twice with ice-cold phosphate buffer saline (PBS), and proteins were extracted using an ice-cold lysis buffer containing 10 mM Tris-HCl (pH 7.5), 50 mM KCl, 2 mM MgCl_2_, 1% Triton X-100, 3 μg/ml aprotinin, 1 μg/ml pepstatin, 1 μg/ml leupeptin, 1 mM dithiothreitol, 0.1 mM Na_3_VO_4_, and 1 mM phenylmethylsulfonyl fluoride. The extracts were incubated on ice for 15 min, then centrifuged at 10,000 × g for 15 min at 4 °C. Supernatants were stored at −80 °C. Protein concentrations were measured using the Bradford assay (Bio-Rad). To evaluate the expression of various proteins, 30 µg of protein extracts from the same set of samples were loaded in parallel onto two identical sodium dodecyl sulfate (SDS)-polyacrylamide gels. After transferring the proteins to Immobilon-P membranes (Millipore), the blots were cut into smaller sections based on the molecular weights of the target proteins. One section was probed for the phosphorylated protein of interest, while the corresponding section was probed for the total protein. The antibodies used for immunoblotting are listed in Supplementary Table 2. Protein visualization was performed using enhanced chemiluminescence (ECL) according to the manufacturer’s instructions (Amersham Biosciences). Band quantification within the linear range of exposure was carried out using ImageJ software.

### Co-immunoprecipitation assay

Cells were rinsed with ice-cold PBS to remove residual media and debris. Cells were lysed using 500 μL of Lysis Buffer A, consisting of 0.5% NP-40, 40 mM HEPES (pH 7.4), 150 mM NaCl, 10% glycerol, 1 μg/mL leupeptin, 2 μg/mL aprotinin, 1 μg/mL pepstatin A, 100 μM PMSF, and Halt phosphatase inhibitor cocktail (Roche, Cat# 11836153001). Lysates were incubated on ice for 5 min with occasional mixing. The soluble fraction was collected by centrifugation at 15,000 rpm for 10 min at 4°C. Protein concentration was determined using a Bradford or BCA assay. Equal amounts of total protein (500-1000 μg) were diluted in 500 μL of Lysis Buffer A. Lysates were incubated with 2 μg of primary antibody at 4°C with gentle rotation overnight. To capture the immunocomplex, 30 μL of Protein G agarose beads or Protein A agarose beads were added, and samples were rotated at 4°C for an additional 2 hours. Beads were collected by centrifugation at 3,000 rcf for 90 seconds, and the supernatant was discarded. Immunoprecipitated proteins were washed three times with Buffer A, with beads collected by centrifugation at 3,000 rcf for 2 min. Samples were subjected to SDS-PAGE and immunoblot analysis as described above.

### Tandem affinity purification of KRAS 4B-G12V binding proteins

FLAG-HIS (FH)-tagged RAS isotypes and KRAS isoforms and their mutants were generated by standard polymerase chain reaction (PCR), subcloned into pCMV2, and transfected into HEK293T cells as previously described (23). Cells were rinsed with PBS and lysed in lysis buffer (0.5% NP-40, 40 mM Hepes (pH 7. 4), 1 mM MgCl₂, 150 mM NaCl, 10% glycerol, 1 mM dithiothreitol (DTT), leupeptin (1 μg/mL), aprotinin (2 μg/mL), pepstatin A (1 μg/mL), and 100 mM 4-(2-aminoethyl)benzenesulfonyl fluoride hydrochloride (AEBSF), Halt phosphatase inhibitor cocktail (Thermo Scientific). The pellet fraction was sonicated in lysis buffer B (40 mM Hepes (pH 7. 4), 1 mM MgCl2, 450 mM NaCl, 10% glycerol, 1 mM DTT, leupeptin (1 μg/mL), aprotinin (2 μg/mL), pepstatin A (1 μg/mL), and 100 mM AEBSF), Halt phosphatase inhibitor cocktail (Thermo Scientific), and proteins were extracted from the 0.5% NP-40 insoluble fraction. Both extracts were combined and subjected to immunoprecipitation using anti-Flag M2-conjugated agarose beads (A2220, Millipore SIGMA) for 2 h at 4°C. The beads were washed six times with wash buffer (WB) (0.5% NP-40, 40 mM Hepes (pH 7.4), 1 mM MgCl2, 150 mM NaCl, 10% glycerol, 1 mM DTT), and the FH-RAS protein complex was eluted with 100 μg/mL 3xFlag peptide (Millipore Sigma). The eluates were subjected to metal affinity purification with Talon bead (Millipore Sigma) for 90 min at 4°C, then washed with WB six times, and FH-RAS complex was eluted with 100 mM EDTA and precipitated with methanol/chloroform for tryptic digestion and LC-MS/MS analysis as described (Sumita et al., Mol Cell 2016).

### Tandem mass spectrometry

Protein samples were reduced with DTT, alkylated with iodoacetamide and digested overnight with trypsin at pH=8.3. Tryptic peptides were extracted and then analyzed by data-dependent reversed-phase microcapillary tandem mass spectrometry (LC-MS/MS) using a hybrid LTQ Orbitrap XL 2D high resolution mass spectrometer (Thermo Fisher Scientific) operated in positive ion mode at a flow rate of 300 nL/min. A 75µm (ID) x 15µm (ID tip) PicoFrit microcapillary column (New Objective, Woburn, MA) was self-packed with Magic C18 resin (Michrom Bioresources, Auburn, CA) to 15 cm (length). The column was equilibrated, and peptides were loaded using buffer A (0.1% formic acid/0.9% acetonitrile/99% water) then eluted with a gradient from 5% buffer B (acetonitrile) to 38% B, followed by 95% B for washing using an EASY-nLCII splitless nanoflow HPLC (Proxeon Biosciences). One MS survey scan was followed by eight MS/MS scans. The Mascot search algorithm (Matrix Science) was used for database searching of all MS/MS spectra against the reversed and concatenated Swissprot all-species protein database with the variable modifications: oxidation (+16 Da) of Met and fixed modifications: carbamidomethyl (+57 Da) of Cys. Mascot search data was incorporated into the Scaffold 3 (Proteome Software) protein identification software with a false discovery rate (FDR) for peptide identifications ∼ 1.5% and less than 1% for protein identifications.

### Expression of eIF2Bα dimers and eIF2B β,γ,δ,ε tetramers labeled with mNeonGreen on eIF2Bβ

Expression vectors pRSF-Duet1 and pCDF-Duet1 (*Novagen*) were used as backbones to construct the three plasmids used to express the eIF2B βγδε tetramers and eIF2Bα dimers. Plasmids PFE-ALTOS-096 (pCDF His8-tev-eIF2Bβ-mNeonGreen/eIF2Bδ) and PFE-ALTOS-026 (pRSF eIF2Bγ/eIF2Bε) were used to coexpress the human eIF2Bβδ and eIF2Bγε protein pairs, respectively. The mNeonGreen fluorescent protein is fused to the C-terminus of subunit eIF2Bβ. Plasmid PFE-ALTOS-041 (pRSF His8-MBP-thrombin-eIF2Bα) was used to express eIF2B𝛂𝛂 fused to an N-terminal His8-Maltose Binding Protein carrier (MBP) to enhance protein expression and stability. TEV and thrombin protease cleavage sites were introduced to remove the purification tags (His8-and His8-MBP-). eIF2Bβγδε tetramers and eIF2Bα dimers were expressed in *E. coli* BL21(DE3)* cells grown in LB media. Cells grew at 37°C until OD600 reached 0.8 before shifting the temperature to 17°C and inducing protein expression with 0.8 mM IPTG for 16 hours. Cells were harvested and washed before processing.

### Purification of eIF2Bα dimers

The His8-MBP-eIF2Bα fusion protein was purified by tandem affinity using Immobilized-Metal Affinity Chromatography (Cobalt-IMAC on Talon resin, *Takara*) followed by amylose-affinity chromatography (Amylose resin, *NEB*). The N-terminal His8-MBP carrier was cleaved using thrombin and eIF2Bα dimers were separated by size exclusion chromatography (SEC) on a Superdex 75 HiLoad 16/60 column (*Cytiva*). The peak corresponding to pure eIF2B𝛂𝛂 dimer was passed through a small column containing 500 𝜇𝜇l of amylose resin and 500 𝜇𝜇l of Cobalt-IMAC resin to subtract any traces of residual His8-MBP or His8-MBP-thrombin-eIF2B𝛂𝛂 proteins.

### Purification of mNeonGreen-labeled eIF2Bβγδε tetramers

Fluorescent eIF2B tetramers were purified by Cobalt-IMAC followed with anion-exchange on a MonoQ HR 10/100 column (*Cytiva*) as described in (66). The peak corresponding to fluorescent tetramers was treated with thermostable hyperTEV (67) to remove the histidine tag from eIF2Bβ-mNeonGreen subunits. Tag-less fluorescent tetramers were then purified by SEC on a Superose 6 HR 10/100 GL (*Cytiva*). The peak corresponding to pure mNeonGreen-labeled tetramers was passed through a small column containing 200 𝜇𝜇l of Cobalt-IMAC resin to subtract any traces of residual histidine-tagged proteins.

eIF2Bα dimers and fluorescent eIF2Bβγδε tetramers were purified and conditioned in the same final SEC buffer composed of 200 mM KCl - 20 mM Hepes pH= 7.2 - 1% glycerol - 5 mM MgCl_2_-0.5 mM TCEP.

### Protein purification for In vitro membrane reconstitution assay

Recombinant proteins were purified as previously described (40, 42, 43, 47, 68) The following proteins were used: human Linker for Activation of T cells (LAT) with a C-terminal hexahistidine tag (residues 27–233); full-length human Grb2; full-length human SOS1; the catalytic domain of SOS1 (SOSCAT, residues 566–1049); and human H-RAS (residues 1–181) with a C118S mutation. Purified LAT was phosphorylated by an ancestrally reconstructed Syk-family kinase domain in the presence of ATP. The kinase was then removed to isolate the phosphorylated LAT (pLAT). A decameric eIF2B complex was assembled by combining a slight molar excess of the eIF2Bα dimer with two mNeonGreen-labeled eIF2Bβγδε monomers and incubating the mixture on ice. A RAS-GTP sensor was engineered by introducing a K65E mutation into the RAS-binding domain of human Raf1 (residues 56–131). This sensor was subsequently purified and labeled with SNAP-Surface Alexa-647 (New England Biolabs) following the manufacturer’s instructions.

### Preparation of supported lipid bilayers

Supported lipid membranes were prepared via the fusion of small unilamellar vesicles (SUVs) onto a Piranha-etched glass substrate, following a previously established protocol. A lipid mixture was prepared by combining 1,2-dioleoyl-sn-glycero-3-phosphocholine (DOPC), L-α-phosphatidylinositol-4,5-bisphosphate (PIP_2_), 1,2-dioleoyl-sn-glycero-3-phosphoethanolamine-N-maleimide (18:1 PE MCC), and 1,2-dioleoyl-sn-glycero-3-[(N-(5-amino-1-carboxypentyl)iminodiacetic acid)succinyl] (nickel salt) (18:1 DGS-NTA) from Avanti Polar Lipids. Lipids were mixed at a molar ratio of 92:2:2:4, respectively, and dried under vacuum to form a thin film totaling 1 μmol. The lipid film was then hydrated with 2 mL of distilled water and sonicated to produce a 1 mM clear lipid solution. A microfluidic channel was assembled using a sticky-Slide well (ibidi, 80608) and a Piranha-etched glass coverslip. The lipid solution was diluted to 0.25 mM in phosphate-buffered saline (PBS) and injected into the channel, followed by the incubation for 30 minutes at room temperature to allow SUVs to spontaneously fuse to the substrate, forming a supported lipid bilayer. Unfused vesicles were washed away with PBS. The channels were then incubated with 100 mM NiCl2 in Tris-buffered saline (TBS) for 5 minutes, followed by a 1 mg/mL bovine serum albumin (BSA) solution. PBS washes were performed between each incubation step.

### Preparation of pLAT/RAS-bound membranes

Immobilization of pLAT onto supported lipid bilayers was achieved by incubating the supported lipid bilayers with 75 nM pLAT solution for 40 minutes, leveraging Ni-NTA chelation via a histidine tag. Unbound protein was subsequently removed with a TBS wash. For H-RAS attachment, supported lipid bilayers were incubated with 24 μM H-RAS solution in TBS supplemented with 5 mM MgCl_2_ for 2.5 hours. This step promoted the thiol-maleimide ligation between the C181 residue of the H-RAS C-terminus and 18:1 PE MCC. The formation of RAS-GDP-bound membranes was completed by a 20-minute incubation with 5 mM β-mercaptoethanol (BME), followed by an overnight incubation with 1 mM GDP at 4 °C. To generate RAS-GTP-bound membranes, the RAS-GDP-bound membranes were treated with 0.5 μM SOSCAT and 1 mM GTP for 30 minutes to ensure complete conversion of RAS-GDP to RAS-GTP. Unbound SOSCAT was then removed with a PBS wash. Before experiments for the analysis of eIF2B membrane recruitment and RAS activation assay, the channels were equilibrated with a buffer containing TBS, 5 mM MgCl2, 1 mM ATP, 10 mM GDP, 0.1 mM orthovanadate (OV), and 0.1 mg/mL BSA.

### Analysis of eIF2B recruitment to membranes

The recruitment of eIF2B to the supported lipid membrane was initiated by injecting a solution containing mNeonGreen-labeled eIF2B into the channels at time zero. The final concentrations of the proteins in the solution were 750 nM Grb2, 5 nM mNeonGreen-eIF2B, and 5 nM SOS. All solutions were prepared in a base buffer containing Tris-buffered saline (TBS), 5 mM MgCl2, 1 mM ATP, 0.1 mM OV, and 0.1 mg/mL BSA. For RAS-GDP-bound membranes, the base buffer included 1 mM GDP, while for RAS-GTP-bound membranes, 1 mM GTP was used. Membrane recruitment of mNeonGreen-eIF2B was monitored using total internal reflection fluorescence microscopy, and the mean fluorescence intensity within a region of interest was measured every 15 seconds.

### Analysis of SOS-mediated RAS activation

The effect of eIF2B on SOS-mediated RAS activation was investigated by injecting a protein solution into the membrane-containing channels at time zero. The final protein concentrations were 750 nM Grb2, 50 nM mNeonGreen-eIF2B, 0.04-1 nM Alexa Fluor 555 (AF555)-labeled SOS, and 10 nM of a RAS-GTP sensor. The buffer used for all solutions included TBS, 5 mM MgCl2, 1 mM ATP, 1 mM GTP, 0.1 mM OV, and 0.1 mg/mL BSA. The mean fluorescence intensities of both the RAS-GTP sensor and AF555-labeled SOS were recorded every 15 seconds for a total of 25 minutes. To normalize the data, the remaining RAS-GDP was fully converted to RAS-GTP by a 25-minute incubation with 560 nM SOSCAT at the end of the experiment. The resulting mean fluorescence intensity that reached equilibrium was then used as a normalization factor to determine the total RAS density and to calculate the percentage of RAS-GTP. The RAS nucleotide exchange rate per SOS molecule (kobs) was calculated based on the assumption of a second-order reaction kinetics. The rate of SOS-catalyzed RAS nucleotide exchange was modeled by the following equation:

dσ[RAS-GTP]/dt =kobs⋅σ[SOS]⋅σ[RAS-GDP]/σ[RAS_total_]

where σ denotes the surface density of the indicated species.

### Synthesis of biotinylated ISRIB

Following a reported procedure (69), to a solution of the known ISRIB analogue (ISRIB, 70 mg, 0.13 mmol, 1.0 equiv. (70)) in DMF (1.3 mL), biotin-PEG4-amine (66.2 mg, 0.143 mmol, 1.1 equiv., commercially available CAS: 663171-32-2), DIPEA (68 μL, 0.39 mmol, 3.0 equiv.), and HATU (59.3 mg, 0.156 mmol, 1.2 equiv.) were added sequentially. The resulting mixture was stirred at room temperature for 10 h, then brine (5 mL) was added, and the mixture was extracted with EtOAc (3 × 15 mL). The combined organic layers were washed with brine, dried over anhydrous Na2SO4, filtered,

and concentrated in vacuo. The residue was purified by silica gel column chromatography (DCM– MeOH gradient elution) to afford CA-P48 as a white solid (79.2 mg, 0.081 mmol, 62%). R*_f_* = 0.28 (10% MeOH in DCM); ^1^H NMR (400 MHz, CDCl_3_) δ 7.43 (d, *J* = 11.5 Hz, 1H), 7.22 (d, *J* = 8.9 Hz, 2H), 6.88 (br, 1H), 6.83 (d, *J* = 9.0 Hz, 2H), 6.80 (d, *J* = 8.5 Hz, 1H), 6.62 (dd, *J* = 8.4, 2.1 Hz, 1H), 6.58 (d, *J* = 8.0 Hz, 1H), 6.50 (d, *J* = 8.3 Hz, 1H), 6.47 (t, *J* = 2.0 Hz, 1H), 6.26 (s, 1H), 5.74 (d, *J* = 10.6 Hz, 1H), 4.76 – 4.67 (m, 1H), 4.48 – 4.43 (m, 1H), 4.39 (s, 2H), 4.30 – 4.22 (m, 1H), 3.91 – 3.70 (m, 4H), 3.63 – 3.45 (m, 19H), 3.41 (d, *J* = 26.4 Hz, 3H), 3.10 (q, *J* = 6.3 Hz, 1H), 2.92 – 2.80 (m, 1H), 2.68 (d, *J* = 12.7 Hz, 1H), 2.17 (t, *J* = 7.2 Hz, 2H), 2.04 – 1.93 (m, 3H), 1.87 – 1.78 (m, 1H), 1.73 – 1.52 (m, 4H), 1.45 – 1.26 (m, 6H) ppm. ^13^C NMR (101 MHz, CDCl_3_) δ 174.3, 170.1, 168.0, 168.0, 167.2, 164.1, 155.9, 141.0, 141.0, 135.3, 129.7, 127.3, 127.1, 118.7, 117.6, 116.1, 112.4, 73.4, 70.7, 70.6, 70.1, 69.8, 69.7, 67.6, 61.9, 60.3, 55.8, 55.2, 49.6, 47.6, 47.3, 40.6, 39.1, 35.8, 31.3, 28.2, 28.1, 25.6 ppm. HPLC (Prodigy^TM^ ODS-3 analytical column, eluent 60 to 95% MeCN:H_2_O/1% TFA buffer, 1.0 mL/min) t_R_ 3.5 min (96%). HRMS: Calc. for C_45_H_63_Cl_2_N_7_O_11_SNa [M+Na]^+^ = 1002.3576 m/z, found [M+Na]^+^ = 1002.3570 m/z.

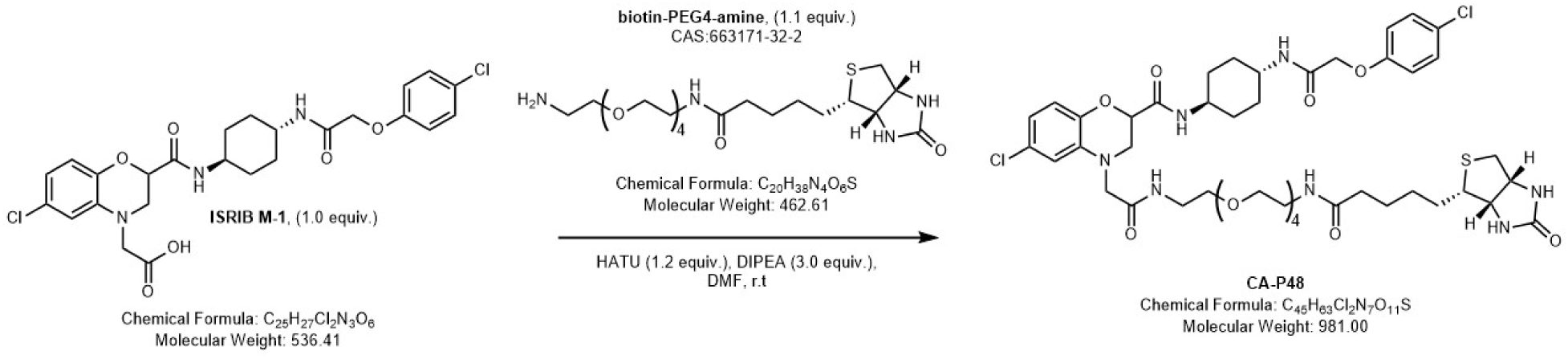

### Biotinylated ISRIB pull-down assay

Cell extracts (500–1000 µg in 500 µL Lysis Buffer A (23)) were incubated with 20 µM biotinylated ISRIB at 4 °C for 30 min with gentle rotation. Separately, 30 µL of streptavidin agarose beads (Thermo Fisher Scientific, Cat# 20347) were washed three times in Lysis Buffer A to remove preservatives, then added to the ISRIB-BIOTIN treated lysates. Mixtures were incubated for 1 hour at 4 °C with gentle rotation. Beads were precipitated by centrifugation at 3,000 × rcf for 90 seconds, and supernatants were discarded. Beads were washed three times with 500 µL Lysis Buffer A and collected with centrifugation at 3,000 × rcf for 2 min each. Bound proteins were eluted by boiling in 2× SDS-PAGE sample buffer for 5 min and analyzed by SDS-PAGE followed by immunoblotting.

### Active RAS pull-down assay

Active RAS was isolated using the active RAS pull-down and detection Kit (Thermo Fisher Scientific, Cat# 16117; Cytoskeleton Inc Cat# BK008) following the manufacturer’s protocol. Briefly, adherent cells were cultured in appropriate growth media and maintained at 70–80% confluence. Cells were washed twice with ice-cold PBS and lysed using the provided 1X Lysis Buffer supplemented with protease inhibitors. The cell lysate was collected by scraping and transferred to microcentrifuge tubes, followed by incubation on ice for 5 min with gentle mixing. Lysates were clarified by centrifugation at 16,000 × g for 15 min at 4°C, and the supernatant was transferred to a new tube. Protein concentration was measured using the BCA assay, and equal amounts of protein (500–1000 µg) were incubated with GST-fusion protein encompassing the RAS binding domain (RBD) of RAF1 bound to Agarose beads for 1 hour at 4°C with gentle rotation. After incubation, beads were precipitated by centrifugation at 10,000 × g for 30 seconds at 4°C and washed three times with ice-cold Lysis Buffer. Bound proteins were eluted by boiling in 2X SDS sample buffer for 5 min at 95°C. The pull-down samples were analyzed by SDS-PAGE and immunoblotting using appropriate antibodies.

### Compound treatment in cell-free assays

AMG-510 (Cat# C-1499) and RMC-6291 (Cat# C-1330) were obtained from ChemGood; BI-3406 (Cat# S8916) and ISRIB (Cat# S0706) from SelleckChem. Each compound was diluted in Lysis Buffer A to the desired concentration and added to freshly prepared extracts in the same buffer (23).. Reactions were incubated for 30 min at 4 °C with gentle rotation. Treated extracts were used directly for active RAS pull-down assays as described above.

### Detection of KRAS:SOS1:eIF2B complex formation with recombinant proteins

To assess ternary complex formation, 2 µg each of HIS-tagged human KRAS G12V (Acro Biosystems, Cat# KRS-H5143), Biotin-FLAG–tagged truncated SOS1 (BPS Bioscience, Cat# 100753), and recombinant tetrameric eIF2B (eIF2Bβγδε; (71)) were combined in 500 µL Lysis Buffer A. ISRIB was treated directly to a final concentration of 5 µM, with an equivalent volume of DMSO used as a negative control. Mixture was incubated at 4 °C for 30 min with gentle rotation. Subsequently, 30 µL of streptavidin-beads were added to each reaction (Sepharose beads served as the tag-negative control) and incubated for 1 h at 4 °C with gentle rotation. Beads were washed three times with Lysis Buffer A and boiled in 2× SDS-PAGE loading buffer for 5 min to elute bound proteins. Complexes were resolved by SDS-PAGE and detected via immunoblotting using antibodies against KRAS, FLAG, and eIF2B subunits.

### Plasma membrane protein isolation and cell fractionation assays

Subcellular fractionation was performed using the Minute Plasma Membrane Protein Isolation and Cell Fractionation Kit (Invent Biotechnologies, Cat# SM-005) according to the manufacturer’s protocol. Briefly, cells were harvested by trypsinization, washed with ice-cold PBS, and lysed in Buffer A. The lysate was filtered and centrifuged at 700 ×g for 1 minute at 4°C to separate the intact nuclear fraction (pellet). The resulting supernatant was centrifuged at 10,000 ×g for 5 min at 4°C to separate the cytosolic fraction (supernatant). The pellet was resuspended in 200 μL of Buffer B, vortexed briefly, and centrifuged at 7,800 ×g for 5 min at 4°C to separate the organelle fraction (pellet). The supernatant was transferred to a new tube, mixed with 1.6 mL of ice-cold PBS by inversion for 30 seconds, and centrifuged at 16,000 ×g for 30 min at 4°C to isolate the plasma membrane fraction (pellet). The intact nuclear, organelle, and plasma membrane fractions were resuspended in RIPA lysis buffer. Protein concentration was determined using a Pierce BCA Protein Assay Kit (Thermo Fisher Scientific, Cat#23227), and each fraction was validated by SDS-PAGE and immunoblotting using appropriate antibodies with distinct cell fraction marker.

### Immunofluorescence staining and confocal imaging

Cells grown on coverslips were washed twice with ice-cold PBS and fixed in 4% paraformaldehyde for 30 min in the dark. After quenching with 50 mM NH₄Cl for 10 min, samples were washed twice in PBS, permeabilized with 0.2% Triton X-100 for 10 min, and blocked for 30 min in PBS containing 2.5% bovine gelatin and 2.5% BSA. Primary antibodies (1:100 in blocking buffer) were applied for 1 h at 4 °C, followed by four PBS washes and incubation with Alexa Fluor– conjugated secondary antibodies (1:500) for 30 min at room temperature. After three additional PBS washes and three rinses in distilled water, coverslips were mounted in Vectashield (Vector Laboratories, Cat# H-1000).

For plasma membrane labeling, cells were washed once in ice-cold HBSS + 0.5% BSA, then incubated with 1 µg/mL Alexa Fluor-conjugated cholera toxin B subunit (CT-B, Invitrogen, Cat# C34778) in the same buffer for 30 min at 4 °C. Cells were washed three times in HBSS + BSA and fixed in 4% paraformaldehyde for 15 min at 4 °C before proceeding with antibody staining as above. Alternatively, CellMask dye (1:5,000 in growth medium) was applied for 1 h prior to fixation.

Immunofluorescence samples were imaged on a Nikon AX/R confocal microscope using NIS-Elements software. Z-stacks were acquired at 0.15 µm intervals, and three-dimensional reconstructions were generated in ImageJ using the 3D Viewer plugin. For colocalization analysis, maximum-intensity projections were converted to 8-bit in ImageJ, and thresholds were set based on control images. Colocalization between CT-B and FLAG-KRAS, or between CellMask and GFP-tagged constructs (GFP-KRAS G12V, GFP-HRAS G12V), was quantified using the Manders’ coefficient plugin (Wright Cell Image Facility).

### Electron Microscopy Imaging and Spatial Statistics

Basal plasma membrane sheets of T47D, H1703, Caco-2, or H358 cells were attached to electron microscopy (EM) gold grids and then fixed in 4% paraformaldehyde/0.1% glutaraldehyde as previously described (72–74). Fixed PM sheets attached to the grids were immunogold-labeled with 4.5 nm gold–conjugated antibodies against GFP (for GFP-tagged proteins), GM3 (TCI Cat#A2582), or SM4 (Sigma Cat#O7139) as previously described (74). After thorough rinsing, EM grids were negatively stained with 0.3% uranyl acetate in 2% methyl cellulose and left to dry overnight before imaging.

Electron micrographs were acquired at 100,000× magnification using JEOL JEM-1400 transmission EM. ImageJ was used to select intact 1 μm² regions of the membrane sheet and assign gold-particle coordinates (72–74). Spatial clustering was quantified by Ripley’s *K*-function, standardized to a 99% confidence interval (48). Values of *L(r)–r* exceeding this interval indicate significant clustering; the maximum value (*L_max_*) was used to summarize clustering extent. Statistical differences between conditions were assessed by bootstrap analysis (48).

### RNA-seq analysis

Total RNA of H358, H1703, HCT116, HK2-8 and MiaPaCa2 cells treated with scrambled siRNAs or eIF2Bε siRNAs (four replicates each) was isolated with Trizol (Thermo Fisher Scientific) and RNA-Seq libraries were prepared following the TruSeq Stranded Total RNA protocol (Illumina) according to the manufacturer’s instructions and 50 base single-end reads were obtained using a HiSeq2500 system in Rapid Mode (Illumina). High-quality reads were then mapped to the hg38 (GRCh38.p12) human genome using the standard HISAT2 pipeline (v2.2.1) (75). Transcript quantitation was performed using Salmon (v0.11.3) (76) and summarized at gene level using the R package tximport (v1.22.0) (77). Gene annotations were assigned using biomaRt (v2.50.3) (78). RNA-seq experiment-wise data were managed and normalized using DESeq2 (v1.44.0) (79). Only protein-coding genes and genes not encoded by Y-chromosome or mitochondrial DNA were considered before TPM normalization and differential expression comparisons. Principal components analysis was conducted using the stats package of base R. Data were filtered to only include genes with 10 or more counts in 2 or more samples. The significance of differential gene expression between vehicle and treatment groups was quantified by one-way ANOVA. Genes with absolute log(FC) > 1 and False Discovery rates (FDRs) < 0.05 were considered differentially expressed. Gene set enrichment analysis (GSEA v4.0.3, Broad Institute) was performed on all genes ranked according to fold change, using the Gene Ontology geneset v5.2 (MSigDB)(80, 81). The number of permutations was 1000 and only sets containing between 15 and 500 genes were retained.

### Ribo-seq data alignment

Ribo-seq data were generated following the protocol described in (50), as detailed in the corresponding GEO entry. Sequenced libraries were trimmed using Cutadapt 3.4 (<u>DOI:10.14806/ej.17.1.200</u>) by removing the constant adapter (AGATCGGAAGAGCAC) from reads without allowing any mismatches. Any reads shorter than six nt and without the adapter were discarded. Trimmed reads were demultiplexed using the sample barcode anchored to the 5′ end of reads without allowing any mismatch in the barcodes. The multiplexed reads were then processed for alignments to the mouse transcriptome using the following criteria: First, low-quality reads and reads shorter than 25 were discarded, then reads were sequentially aligned to cytoplasmic and mitochondrion rRNA and tRNA using Bowtie 2 version 2.5.4 (PMID: 22388286) allowing three mismatches in the reads (parameters: −v 3 −norc −p 12). The remaining reads were aligned with the human GRCh38 whole transcriptome preset by Bowtie 2 version 2.5.4. SAM files were converted to BAM files, sorted, indexed.

### Differential expression analysis of Ribo-seq data

Use htseq-count software (82) and set the CDS mode to calculate the number of reads. Differential analysis was performed using the R package deltaTE (ΔTE) version 1.34.0, (51) which incorporates an interaction model into the DESeq2 method. For each gene, a p-adjusted value (p-adj) was calculated for the fold changes between two experimental conditions at Ribo-seq, RNA-seq, and TE levels, and p-adj below 0.05 was considered significant for each fold change. These fold changes were then used to divide genes into eight categories described in the text. For scatterplots displaying the fold change between Ribo-seq and RNA-seq in **Figure 5c**, genes with the p-adj value of “NA” in any fold change (Ribo-seq, RNA-seq, and TE) comparison were excluded from the graph. The values for the PCA plots in **Figure 5b** were also determined using DESeq2. The gene expressions shown in Figure **5e** are generated from raw counts using the variance stabilizing transformation (VST) in DESeq2 (79).

### Enrichment Analysis of Ribo-seq data

Significantly upregulated genes (adjusted *p* < 0.05, |log₂FC| > 0.4) were selected for each defined region. The selected genes were submitted to Metascape (83) for functional enrichment. The analysis was performed using default settings, including GO Biological Processes, KEGG, Reactome, and Hallmark gene sets. Enriched terms were filtered by *p* < 0.01, a minimum count of 3 genes, and an enrichment factor > 1.5. Redundant terms were clustered, and the top term from each cluster was retained.

### Construction of protein‒protein Interaction (PPI) Network

The STRING platform (http://string-db.org) was used to analyze the PPI network (84). A minimum required interaction score > 0.4 (the medium confidence) was set as the cut-off criterium, along with “Homo sapiens” as the organism. Network visualization of core clusters was conducted via utilizing Cytoscape 3.10.1 (85), an open-source software for analyzing and visualizing networks. The MCODE and CytoHubba plug-ins of the Cytoscape 3.10.1 were, respectively, utilized to screen out the core modules. Their default parameters were as follows (for the MCODE plug-in: degree cutoff ≥ 10, node score cutoff ≥ 0.2, K-core ≥ 5, and max depth = 100; for the CytoHubba plug-in: the top 10 nodes chosen by MCC).

### In silico modeling of eIF2B:SOS:KRAS complex

The prediction of the structural assembly of KRAS:SOS:eIF2B was conducted by combining AlphaFold 3 predictions and existing structural information (namely PDB IDs: 7L7G (66), 7KFZ (56), 6W4E (86). A more detailed illustration and explanations on model components is provided in **Suppl Figure 3**. AlphaFold 3 predictions were generated using Google DeepMind’s AlphaFold Server (33). For the prediction of KRAS(G12V):SOS:eIF2B complex, the following UniProt (87) sequences were used: Q13144; P01116-2 (including only residues 1–185 and introducing the G12V mutation); Q07889 (including only residues 566-1046); P49770; Q9NR50; Q9UI10.

The validation information for AlphaFold 3 predictions is provided in **Suppl Figure 9**. For the prediction of KRAS G12V:eIF2Bε subunit complex we used Q13144, P01116-2 (including only residues 1–169 and introducing the G12V mutation), GTP – Guanosine-5’-triphosphate (ligand) and Mg^2+^ (ion) as input. The visualizations and alignment of the structures were done with PyMOL (The PyMOL Molecular Graphics System, Version 3.0 Schrödinger, LLC.)

### TCGA data analysis

The correlation between eIF2B expression and overall patient survival was analyzed using the UCSC Xena online tool (https://xenabrowser.net/). Data sets from TCGA Lung Adenocarcinoma (LUAD) and TCGA Pan-Cancer (PANCAN) cohorts were utilized. Samples containing missense KRAS mutations were identified and filtered using the somatic mutation (SNP and Indel) MC3 Public version data set. Expression levels of eIF2B subunits were stratified into quartiles for comparative analysis.

### Transgenic mouse model and treatment

To impair eIF2Bε in mouse LUAD tumors, KRAS^+/LSL-G12C^ mice from The Jackson Laboratory (Strain #: 033068) were crossed with fTg/0;eIF2α^S/S^ mice (21) and eIF2Bε^f/f^ mice obtained from European Mouse Mutant Archive (EMMA) on a C57BL/6 background. Lung tumorigenesis was induced in KRAS^+/LSL-G12C^;fTg/0;eIF2α^S/S^/eIF2Bε^+/+^ and KRAS^+/LSL-G12C^;fTg/0;eIF2α^S/S^/eIF2Bε^f/+^ mice through intratracheal intubation with lentiviruses expressing CRE and TP53 shRNA (21, 88). Lung tumor development was monitored using ultrasound imaging on the Visual Sonics VEVO 3100 high-frequency ultrasound system (89).

### Xenograft tumor assays

Tumor transplantation assays were conducted in 8-week-old female athymic nude mice (Charles River Inc.) following the established protocol (21). Briefly, cells were suspended in a 1:1 v/v mixture of phosphate-buffered saline (PBS): Matrigel (Corning) and injected subcutaneously into the flanks of nude mice. Mice were inoculated with 5×10^5^ cells in 200 µl PBS: Matrigel mix. Tumor growth was monitored twice per week using digital calipers, and the volume was calculated using the formula: tumor volume [mm^3^] = π/6 x (length [mm]) x (width [mm]) x (height [mm]).

### Immunohistochemistry

Immunohistochemistry was performed at the Segal Cancer Centre Research Pathology Facility (Jewish General Hospital, Montreal). Paraffin-embedded tissue sections (4 µm) were mounted on TOMO slides (VWR) and dried overnight at 37 °C. All staining steps were performed on a Ventana Discovery XT Autostainer using reagents from Roche/Ventana unless stated otherwise. Slides were deparaffinized and subjected to heat-induced epitope retrieval in CC1 solution (prediluted, Cat #950-124) under the standard protocol. Primary antibodies—rabbit polyclonal anti-eIF2Bε (ProteinTech, 1:500), rabbit monoclonal anti-phospho-ERK (Cell Signaling, 1:100), and rabbit monoclonal anti-Ki-67 (Abcam, 1:500)—were diluted in Ventana Antibody Diluent (Cat #251-018) and applied for 32 min at 37 °C. Detection was carried out with the OmniMap anti-Rabbit HRP detection kit (Cat #760-4311) for 8 min, followed by ChromoMap DAB substrate (Cat #760-159). Negative controls omitted the primary antibody. Slides were counterstained with Hematoxylin II (Cat #760-2021) for 12 min, blued with Ventana Bluing Reagent (Cat #760-2037) for 4 min, then removed from the autostainer, rinsed in warm soapy water, dehydrated through graded ethanol, cleared in xylene, and coverslipped with Eukitt Mounting Medium (EMS, Cat #15320). Stained sections were examined by conventional light microscopy or scanned using an Aperio AT Turbo Scanner (Leica Biosystems). For IHC images, the percentage of cells positive for eIF2Bε, ERK-P, Ki-67, and TTF1 was analyzed using QuPath digital pathology software (v0.5.1). Cell types were classified using the cell detection annotation tool in QuPath. Tumor cells were identified based on H&E staining and QuPath’s nucleus-to-cell area ratio analysis, using a threshold value of 0.68(90).

### Guidelines of ethical conduct in mouse work

The animal studies were performed in accordance with the Institutional Animal Care and Use Committee (IACUC) of McGill University and procedures were approved by the Animal Welfare Committee of McGill University (protocol JGH-10070).

### Statistical analysis

Statistical analysis was conducted using three biological replicates. Error bars represent the standard error as indicated, and significance in differences between arrays of data was determined using a two-tailed Student’s t-test.

---

**Suppl Figure 1. eIF2B-mediated stimulation of cell survival and MAPK signaling is selective for tumor cells harboring mutant KRAS.**

Evaluation of the colony forming efficacy and MAPK signaling in:

a. H358 (KRAS G12C) and H1703 (wild-type KRAS) cells after eIF2Bε KD via 2 different shRNAs.
b. H1703 cells stably expressing either GFP-tagged wild-type KRAS or GFP-tagged KRAS G12C.
c. Isogenic HCT116 (KRAS G13D) and HK2-8 (wild-type KRAS) cells following eIF2Bε KD vias siRNAs.

(d) Mouse lung tumor cells expressing KRAS G12D either with wild type eIF2α (eIF2α^S/S^) or non-phosphorylatable eIF2α S51A (eIF2α^A/A^) before and after eIF2Bε KD by siRNAs. Because eIF2α^A/A^ cells exhibited substantially slower growth than eIF2α^S/S^ cells (21), extracts from the former were prepared three days later to obtain sufficient material for immunoblot analysis.

(**a**–**d**) SCR, scrambled siRNA or shRNA used as a negative control. Protein lysates were analyzed by immunoblotting for phosphorylated MEK and ERK. Ratios of phosphorylated to total protein are

indicated. Representative results from three independent experiments are shown. Graphs show quantification from three independent biological replicates; each performed in triplicate. Data are presented as mean ± SEM.

***Suppl Figure 2.* eIF2B interacts preferentially with SOS in the presence of mutant KRAS rather than HRAS or NRAS.**

(a) Interaction of endogenous eIF2Bε, SOS1, and KRAS in H358 cells. Cell lysates from H358 cells were subjected to co-IP using antibodies against the α, β, δ, or ε subunits of eIF2B, as well as SOS1, followed by immunoblotting for SOS1, eIF2Bε, and KRAS. The antibodies used for IP were as follows: eIF2Bα (Proteintech, Cat# 18010-1-AP), eIF2Bβ (Proteintech, Cat# 11034-1-AP), eIF2Bδ #1 (Fortis Life Sciences, Cat# A302-982A-T), eIF2Bδ #2 (Fortis Life Sciences, Cat# A302-983A-T), eIF2Bε #1 (Fortis Life Sciences, Cat# A302-556A-T), eIF2Bε #2 (Fortis Life Sciences, Cat# A302-557A-T), SOS1 #1 (Proteintech, Cat# 55041-1-AP), and SOS1 #2 (Fortis Life Sciences, Cat# A301-890A-T). As a negative control, equal amounts of cell lysate were IPed using non-specific rabbit IgG. Protein loading and expression levels in the co-IP assays were verified by immunoblotting of whole-cell extracts (WCE).

(**b**, **c**) HEK293T cells were transfected with various FLAG-KRAS constructs, including wild-type KRAS (WT), KRAS G12V, and KRAS G12V variants containing C→S and/or polyK→polyQ mutations. In (panel b), cell lysates were subjected to IP using an anti-FLAG antibody, followed by immunoblotting to detect co-IPed endogenous eIF2B subunits. In (panel c), HEK293T cells were co-transfected with MYC-tagged constructs for all eIF2B subunits and the indicated FLAG-tagged KRAS constructs (WT, G12V, or G12V with C→S and polyK→polyQ substitutions). Cell lysates were incubated with ISRIB–biotin, followed by streptavidin pull-down and immunoblotting with anti-FLAG and anti-MYC antibodies.

(**d**) Reconstitution of SOS:eIF2B:KRAS complex with recombinant proteins. A cocktail of purified recombinant proteins, consisting of 2 μg each of biotinylated-FLAG-tagged SOS^CAT^ (residues 566– 1046), eIF2B subunits (β, γ, δ, and ε), and HIS-tagged KRAS G12V, was incubated in the absence or presence of 5 μM ISRIB. Proteins were subjected to pull down assays with streptavidin-Sepharose beads. As a negative control, an equivalent amount of Sepharose beads was added to parallel reaction. Precipitated proteins were subjected to immunoblotting for eIF2B subunits, KRAS or FLAG to detect SOS^CAT^.

(**e**) Complex formation between eIF2B, mutant KRAS, and SOS. HEK293T cells were co-transfected with MYC-tagged constructs for all eIF2B subunits, HA-SOS1 and FLAG-tagged WT KRAS, KRAS G12V, or KRAS G12V containing the C→S and polyK→polyQ mutations. Cell lysates were subjected to IP with an anti-MYC antibody, followed by immunoblotting with anti-FLAG, anti-HA, and anti-MYC antibodies to detect KRAS, SOS1, and eIF2B subunits, respectively, in the co-IPed proteins. Immunoblotting for phosphorylated ERK (ERK-P) in WCE was a readout of FLAG-KRAS activity.

(**f**, **g**) eIF2Bε preferentially interacts with mutant KRAS 4B compared to KRAS 4A, HRAS, or NRAS, in either their wild-type or mutant forms. HEK293T cells were co-transfected with MYC-tagged eIF2Bε and HA-tagged SOS1 together with various FLAG-tagged constructs encoding wild-type HRAS, wild-type KRAS, or their G12V mutant forms (panel f). Similarly, HEK293 cells were transfected with FLAG-tagged variants of HRAS, NRAS, KRAS 4A, and KRAS 4B carrying the G12V mutation (panel g). Cell lysates were subjected to IP using an anti-FLAG antibody, followed by immunoblotting with anti-MYC and anti-HA antibodies to detect eIF2Bε and SOS1, respectively. Protein loading in the co-IP assays was verified by immunoblotting of whole-cell extracts (WCE).

(h) eIF2Bε KD reduces GTP-bound KRAS in H358 cells harboring KRAS G12C. Cell lysates from H358 cells transduced with either scrambled control or eIF2Bε shRNA were subjected to pull-down assays using GST-tagged RBD of RAF, followed by immunoblotting with antibodies against KRAS, eIF2Bε, and GST.

(i) Activation of eIF2B GEF by ISRIB enhances the formation of GTP-bound KRAS and its association with eIF2B and SOS1. H358 cell lysates treated with increasing concentrations of ISRIB were subjected to pull-down using GST-RBD of RAF, and bound proteins were analyzed by immunoblotting for KRAS, SOS1, and eIF2Bα to ε subunits.

(**j**, **k**) ISRIB counteracts the reduction of GTP-bound KRAS induced by SOS inhibition or KRAS G12C inhibition in H358 cells. In panel (j), lysates were treated with the SOS inhibitor BI-3406 and ISRIB; in panel (k), lysates were treated with the KRAS G12C inhibitor AMG510 and ISRIB. Samples were subjected to GST-RBD pull-down assays, followed by immunoblotting to detect KRAS, SOS, and eIF2Bα to ε subunits.

**Suppl Figure 3: *Computational construction of the KRAS:SOS:eIF2B complex*.**

The full model was generated from two AlphaFold 3 (AF3) predictions that independently identified putative interactions between eIF2B and RAS/SOS. Specifically, these AF3-predicted assemblies were the KRAS G12V:SOS^CAT^:eIF2B (ε, β, δ, γ) complex, and the KRAS G12V:eIF2Bε complex. In the first AF3 prediction, KRAS G12V is positioned at the catalytic site of SOS, while the eIF2Bε subunit associates with the allosteric RAS-binding site. In the second model, GTP-bound KRAS G12V interacts with eIF2Bε near the predicted SOS-association region. The validation data of AF3 predictions is available in Suppl Figure 9.

To construct the complete model and validate AF3-derived configurations, existing structural data were incorporated (PDB IDs: 7KFZ, 7L7G, and 6W4E). Alignment of the RAS:SOS module from the first AF3 model with the cryo-EM structure of SOS1:KRAS (PDB 7KFZ;(56)) showed strong agreement. The organization of the eIF2B(ε, β, δ, γ) subcomplex in the AF3 model also closely matched the cryo-EM structure of eIF2B bound to ISRIB (PDB 7L7G;(66)).

As the second AF3 prediction indicated that GTP-bound KRAS could interact with eIF2Bε via its effector lobe, this interaction would be incompatible with the allosteric RAS site of the RAS:SOS:RAS complex. However, the predicted dimerization interface of the allosteric RAS remained accessible and was found to be compatible with the NMR-derived structure of a GTP-bound KRAS dimer in a lipid nanodisc (PDB 6W4E;(86). Based on this, we propose that eIF2Bε can interact with GTP-bound KRAS dimers (or higher-order oligomers) that expose the effector lobe.

Specific parts of the AF3-predicted models that were absent in experimental structures (disordered or missing residues) were omitted from the final model, as they were not predicted to contribute to the key interfaces. In the resulting putative KRAS:SOS:eIF2B assembly, the KRAS HVRs are appropriately oriented toward the membrane, and no steric clashes are apparent among the complex components. The membrane model, which was used for visualization purpose only obtained from a previous study (91).

**Suppl Figure 4. eIF2Bε co-localizes with SOS and mutant KRAS at the PM.**

(**a**, **b**) 3D imaging constructed from Z-stack immunofluorescence of the cells shown in Figure 4a reveals the spatial distribution of MYC-eIF2Bε (red), HA-SOS1 (blue) and FLAG-KRAS (green) either with the G12V mutation (panel a) or in the wild type from (panel b).

(c) Immunoblot analysis of phosphorylated and total ERK in protein extracts from HEK293 cells co-expressing either FLAG-tagged wild-type KRAS or KRAS G12V with MYC-tagged wild-type eIF2Bε, the hyperactive D154A mutant, or the catalytically inactive N263K mutant.

(d) IF analysis of each MYC-eIF2B subunit separately (red), HA-SOS1 (blue) and FLAG-KRAS G12V (green), co-expressed in HEK293T cells. The white signal in the merged images indicates the co-localization of the three proteins at the periphery of the cell.

**Suppl Figure 5. *eIF2B acts as a co-GEF with SOS for RAS in an in vitro SOS-mediated RAS activation assay*.** (**a**) Schematic representation of the physical mechanism of SOS activation. Full-length SOS is autoinhibited in the cytosol and becomes activated upon recruitment to the plasma membrane. When GRB2 binds to phosphotyrosines on LAT, autoinhibited SOS is recruited to Grb2 through an interaction between the SOS proline-rich (PR) domain and the SH3 domain of GRB2. Once localized at the membrane, autoinhibition is relieved following PIP_2_ binding to the autoinhibitory PH domain of SOS, which exposes the allosteric RAS-binding site. Subsequent binding of RAS-GTP to this allosteric site further enhances SOS activation. The probability of SOS activation increases with its membrane dwell time. Once activated, a single SOS molecule is highly processive, catalyzing nucleotide exchange for hundreds of RAS molecules during a single membrane-binding event. Newly generated RAS-GTP reinforces this activation through a positive feedback loop by binding to the SOS allosteric site. (**b**) Schematic of the reconstituted membrane system used to quantitatively analyze eIF2B recruitment and SOS activity. The assay utilizes mNeonGreen-labeled eIF2B and an Alexa Fluor 647 (AF647)-labeled RAS–GTP sensor, allowing for the stepwise analysis of (**c**) eIF2B recruitment to the membrane, (**d**) conversion efficiency of RAS–GDP to RAS–GTP, and (e) the nucleotide exchange rate per SOS molecule.

***Suppl Figure 6.* eIF2Bε predicts poor patient survival with mutant KRAS-driven cancers.**

(**a**, **b**) Kaplan–Meier overall survival analyses reveal that eIF2Bε mRNA expression is significantly associated with patient outcomes in mutant KRAS-driven cancers. In both lung adenocarcinoma (LUAD; panel a) and pan-cancer (PANCAN; panel b) cohorts with *KRAS* missense mutations, elevated eIF2Bε mRNA levels correlate with poor survival. By contrast, in tumors lacking *KRAS* mutations, eIF2Bε expression has minimal impact on overall survival.

**Suppl Figure 7. Prognostic assessment of eIF2B subunit expression in LUAD patient survival.** Analysis of the other eIF2B subunits (α, γ, δ) revealed no significant correlation with patient survival in LUAD datasets, irrespective of *KRAS* mutational status (left panels, wild-type *KRAS*; right panels, *KRAS* missense mutations). An exception is the eIF2Bβ subunit, where low mRNA expression is associated with reduced survival probability in LUAD tumors with wild-type *KRAS*; however, this observation is based on a limited number of cases (n<15).

**Suppl Figure 8. Assessment of the prognostic significance of eIF2B subunit expression in all cancers.** Expression analysis of the other eIF2B subunits (α, γ, δ) across TCGA pan-cancer datasets revealed no significant association with patient survival excluding LUAD patients, irrespective of *KRAS* mutational status (left panels, wild-type *KRAS*; right panels, *KRAS* missense mutations). An exception is the eIF2Bβ subunit, where increased mRNA expression is associated with reduced survival probability across all cancers harboring mutant *KRAS*.

**Suppl Figure 9. Confidence and accuracy of the AlphaFold 3 predicted models.**

(a) KRAS G12V:SOS^CAT^:eIF2B(ε, β, δ, γ) complex. Predicted local distance difference test (pLDDT). (Top) The AlphaFold 3 predicted model of KRAS(G12V):SOS^CAT^:eIF2B(ε, β, δ, γ) complex colored by pLDDT on the right. The model with the colored subunits is given as a refence in the same orientation on the left. The pLDDT indicates reasonable confidence for the different subunits, and near the predicted interaction interfaces. The predicted template modeling (pTM) and the interface predicted template modeling (ipTM) scores are also shown in the image. (Bottom) Predicted aligned error matrix. The magenta rectangle in the matrix highlights the parts of eIF2Bε and SOS^CAT^ located in their predicted interaction interface, which are shown in the 3D-structure in green and orange, respectively.

(b) <u>KRAS(G12V):eIF2Bε complex</u>. (Top) Predicted local distance difference test (pLDDT). The AlphaFold 3 predicted model of KRAS(G12V):eIF2Bε complex colored by pLDDT on the right. The model with the colored subunits is given as a refence in the same orientation on the left. The pLDDT indicates reasonable confidence for the different subunits, and near the predicted interaction interfaces. The predicted template modeling (pTM) and the interface predicted template modeling (ipTM) scores are also shown in the image. (Bottom) Predicted aligned error matrix.

