## Supplementary material for "eIF2B Selectively Anchors and Activates Mutant KRAS": Suppl Figures 1_9

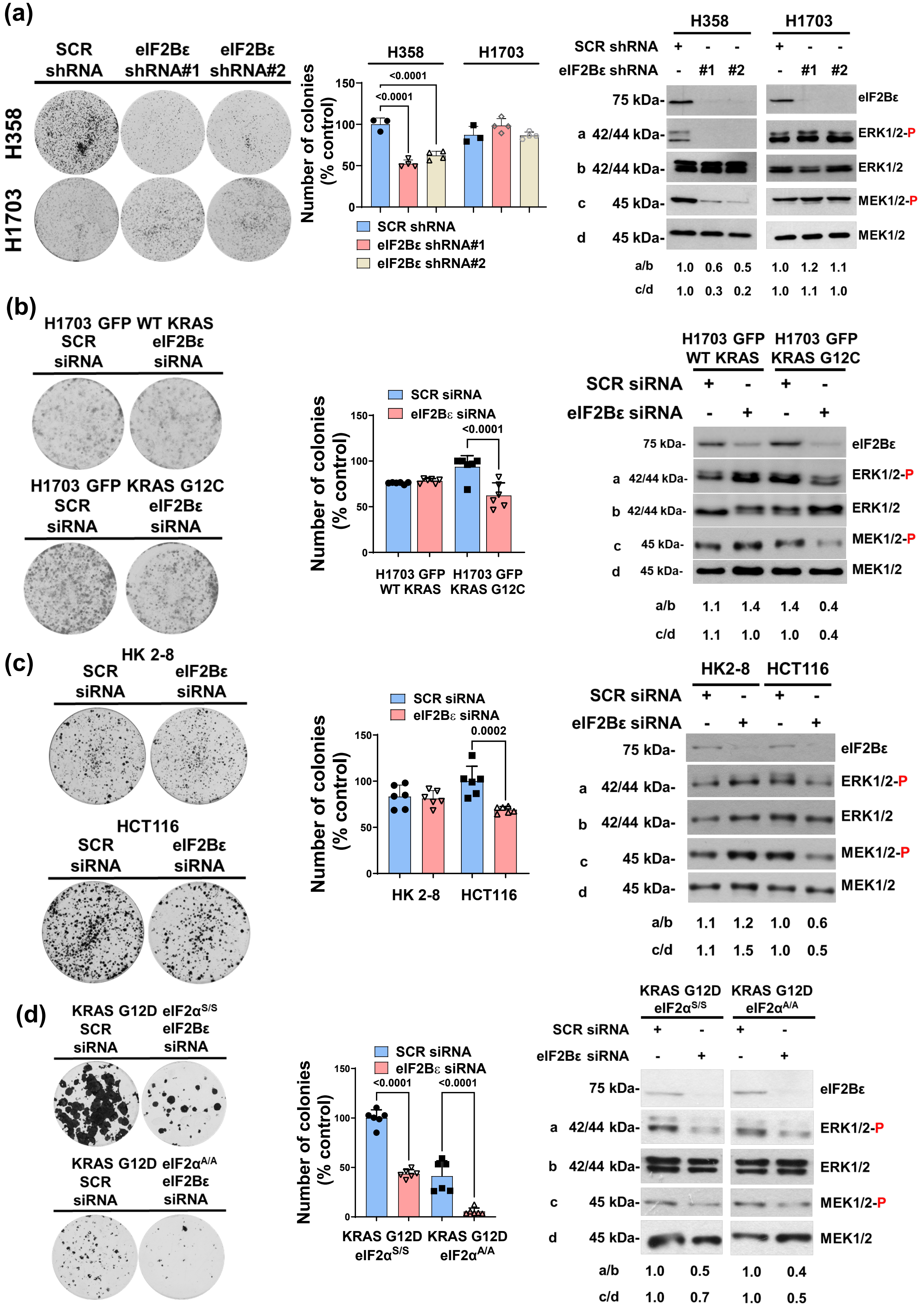


**Suppl Figure 1**

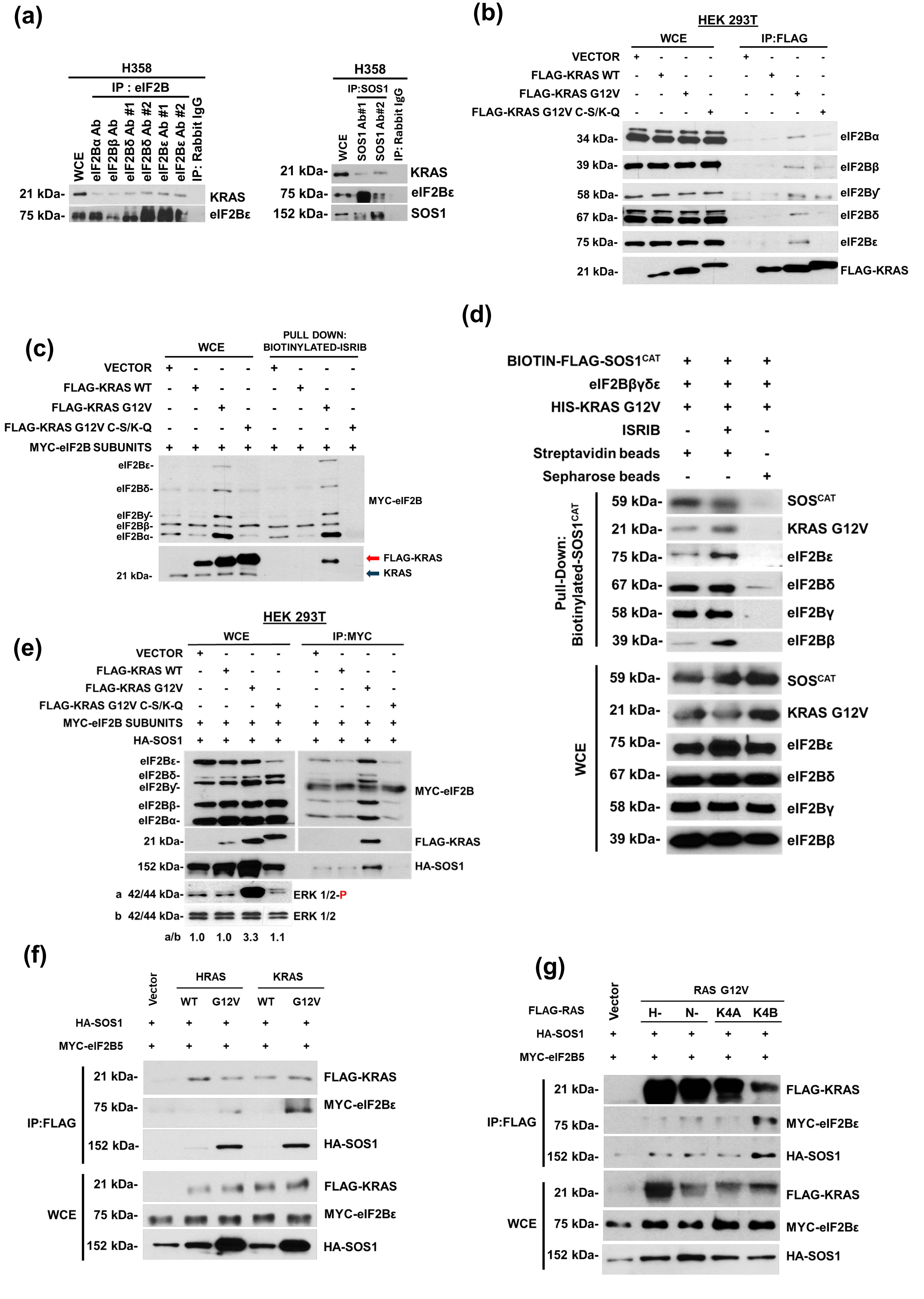


Suppl Figure 2

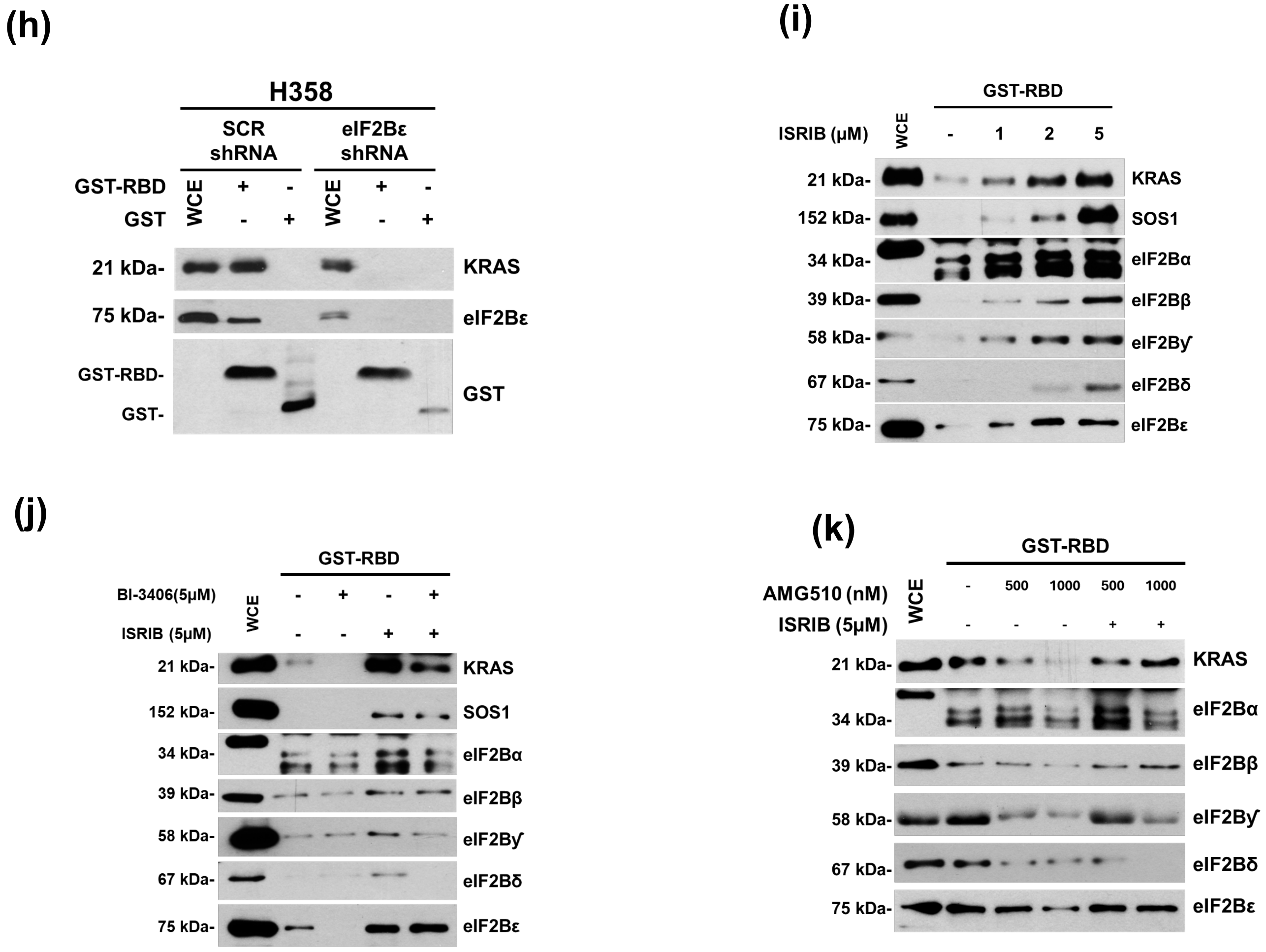


**Suppl Figure 2 (cont'd)**

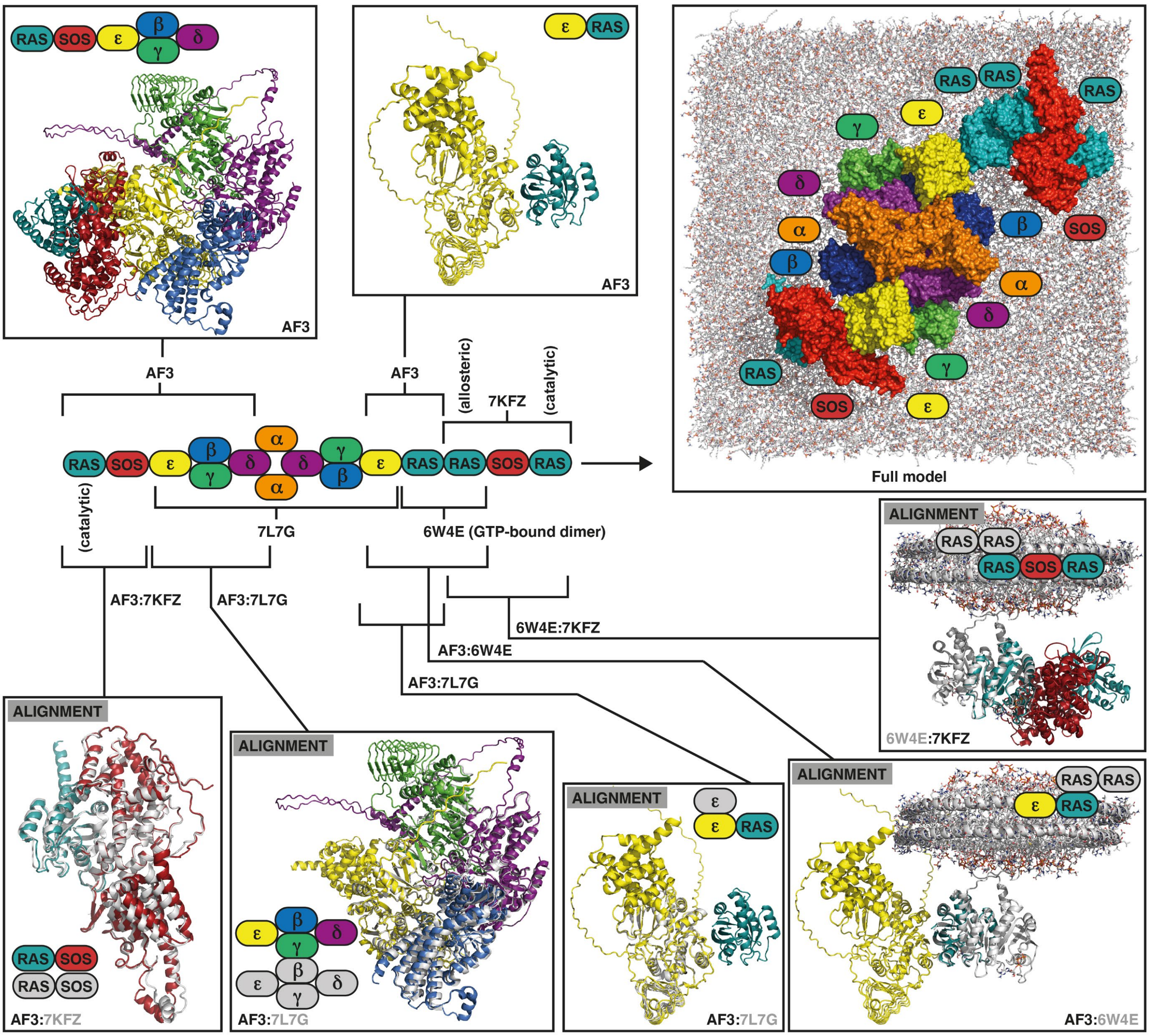


Suppl Figure 3

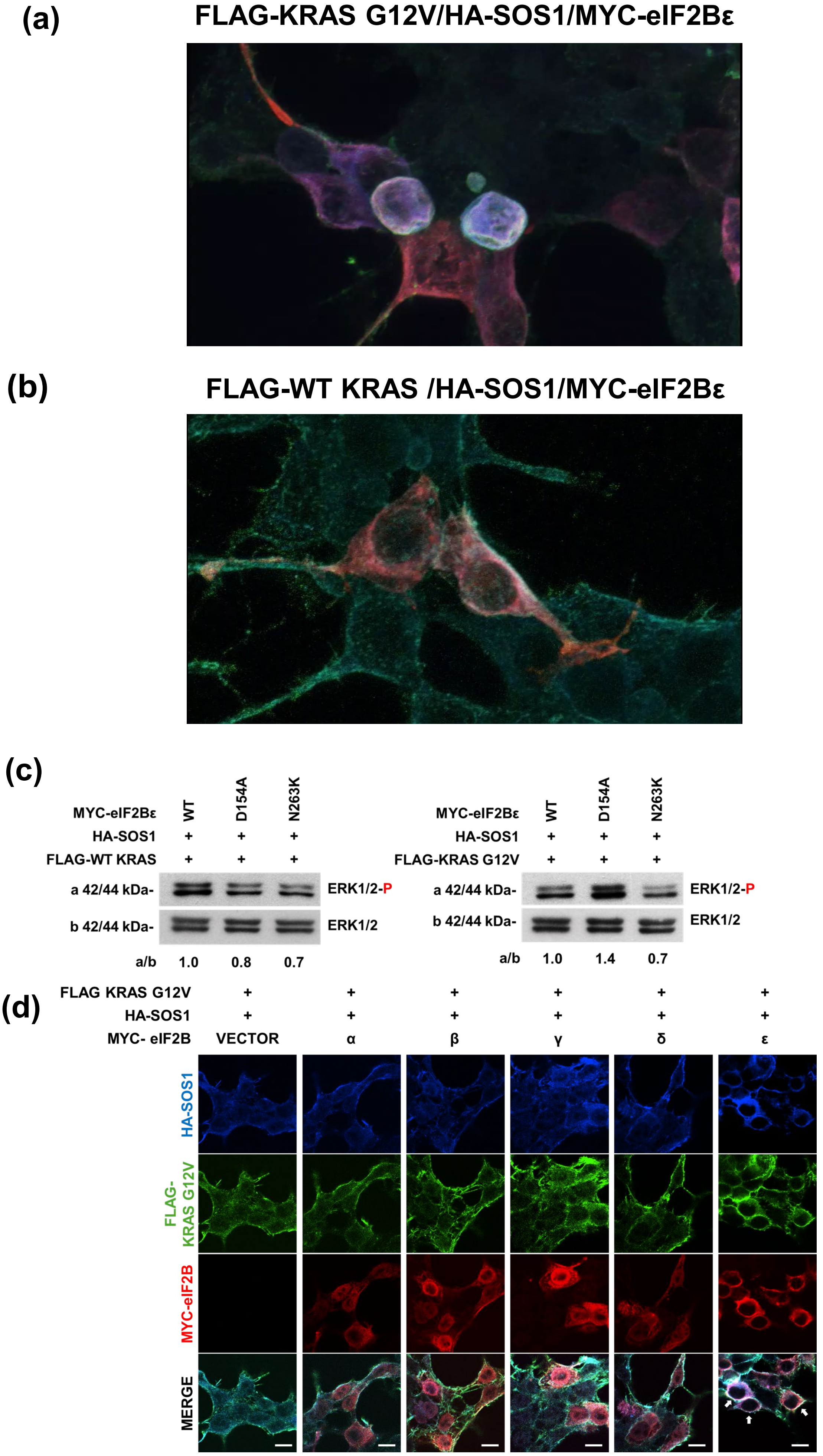


Suppl Figure 4

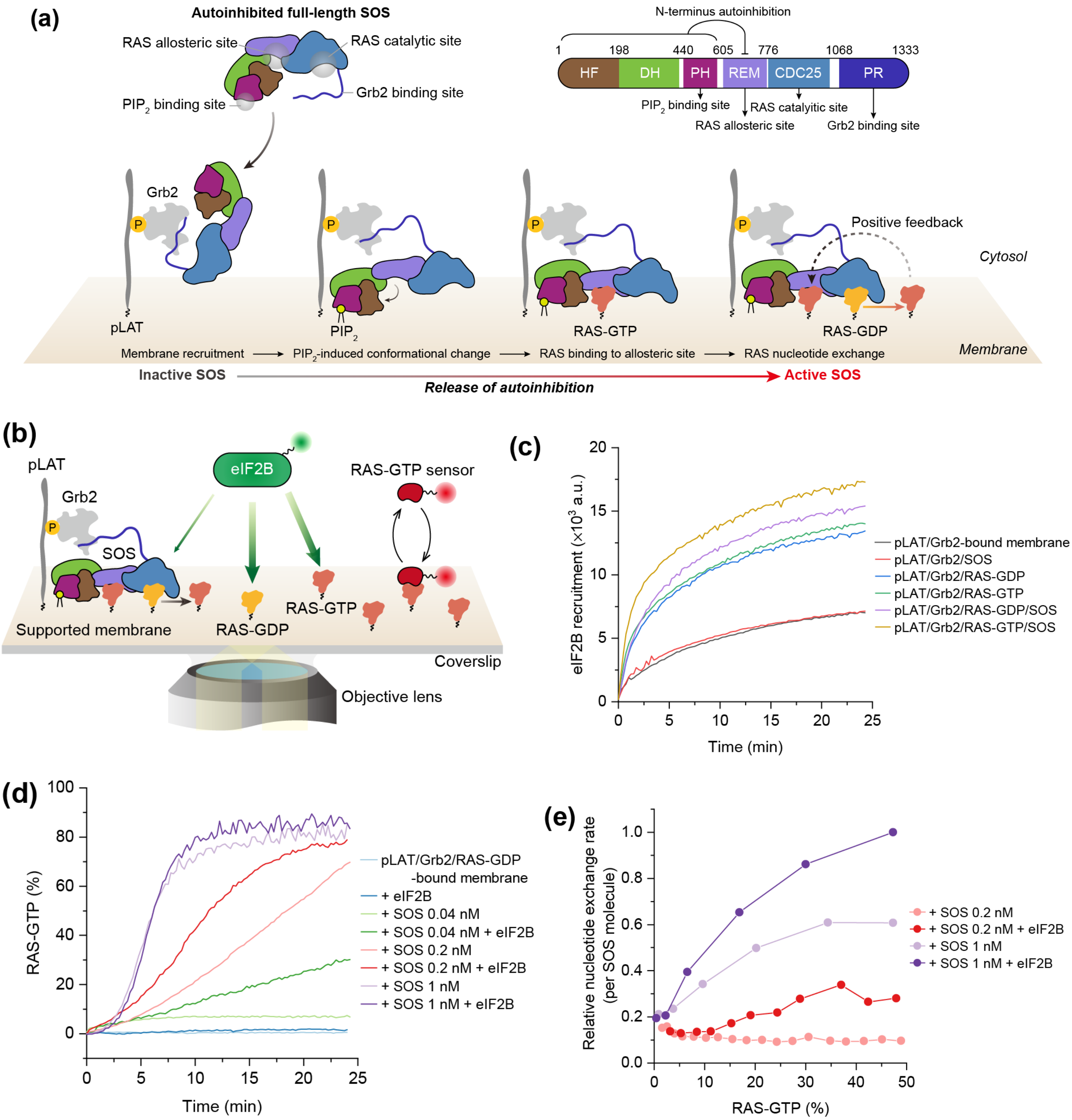


### Suppl Figure 5

**(a)**

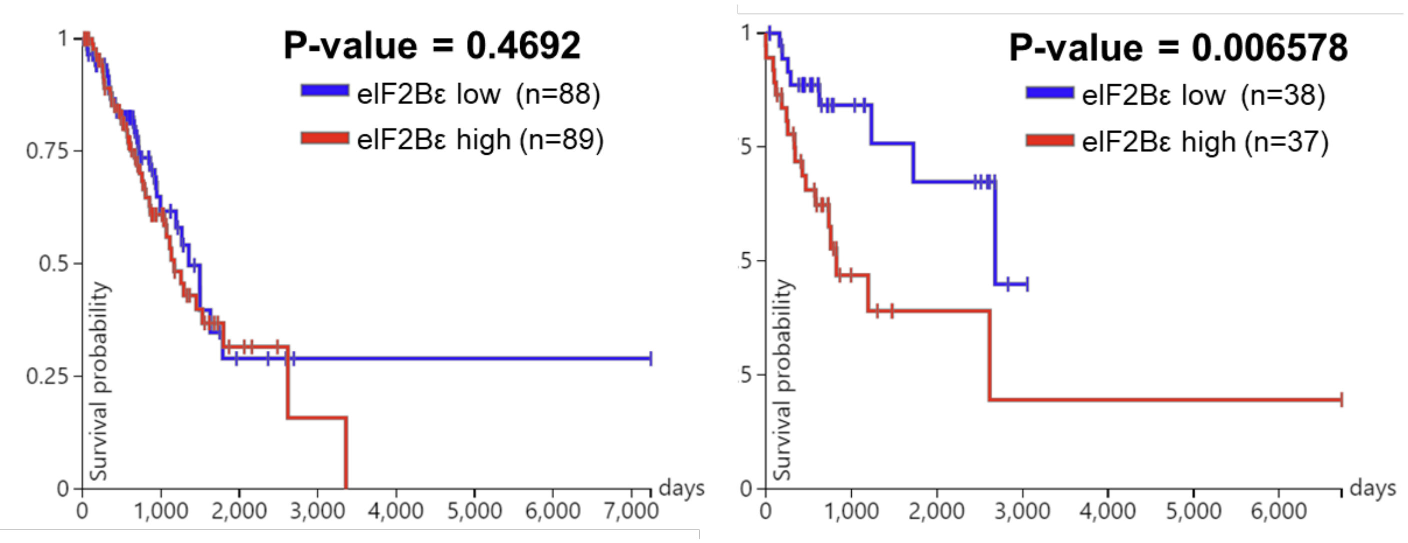


**TCGA LUAD**
**Wild-Type KRAS**

**TCGA LUAD**
**Mutant KRAS**

**(b)**

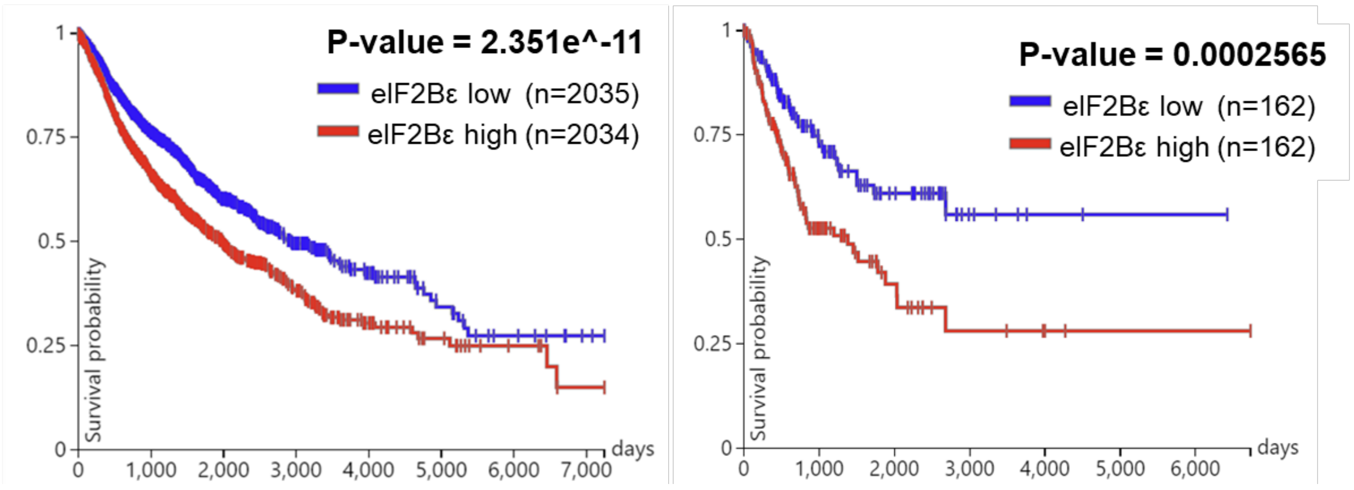


**TCGA Pan-CAN**
**Wild-Type KRAS**

**TCGA Pan-CAN**
**Mutant KRAS**

**Suppl Figure 6**

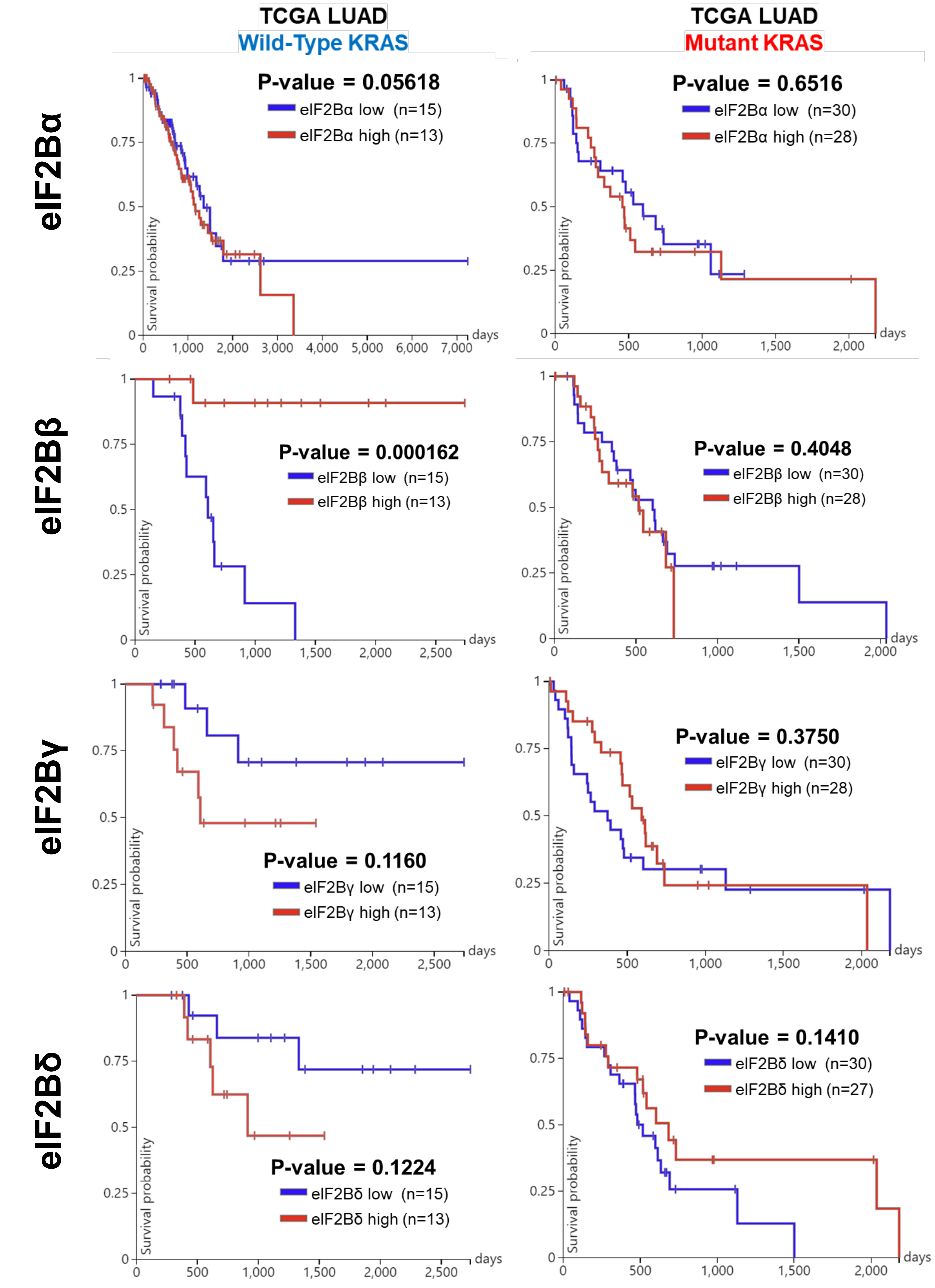


Suppl Figure 7

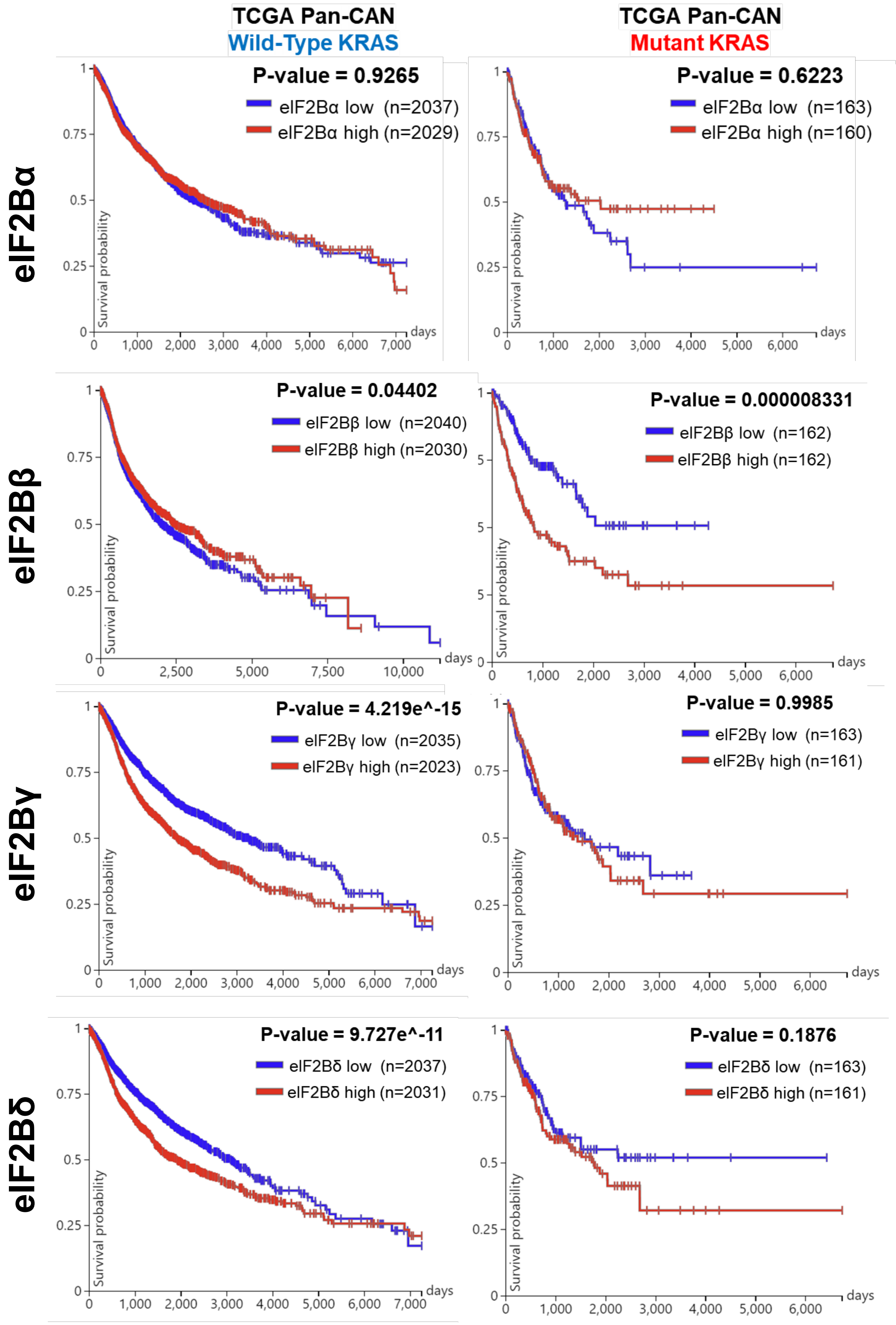


Suppl Figure 8

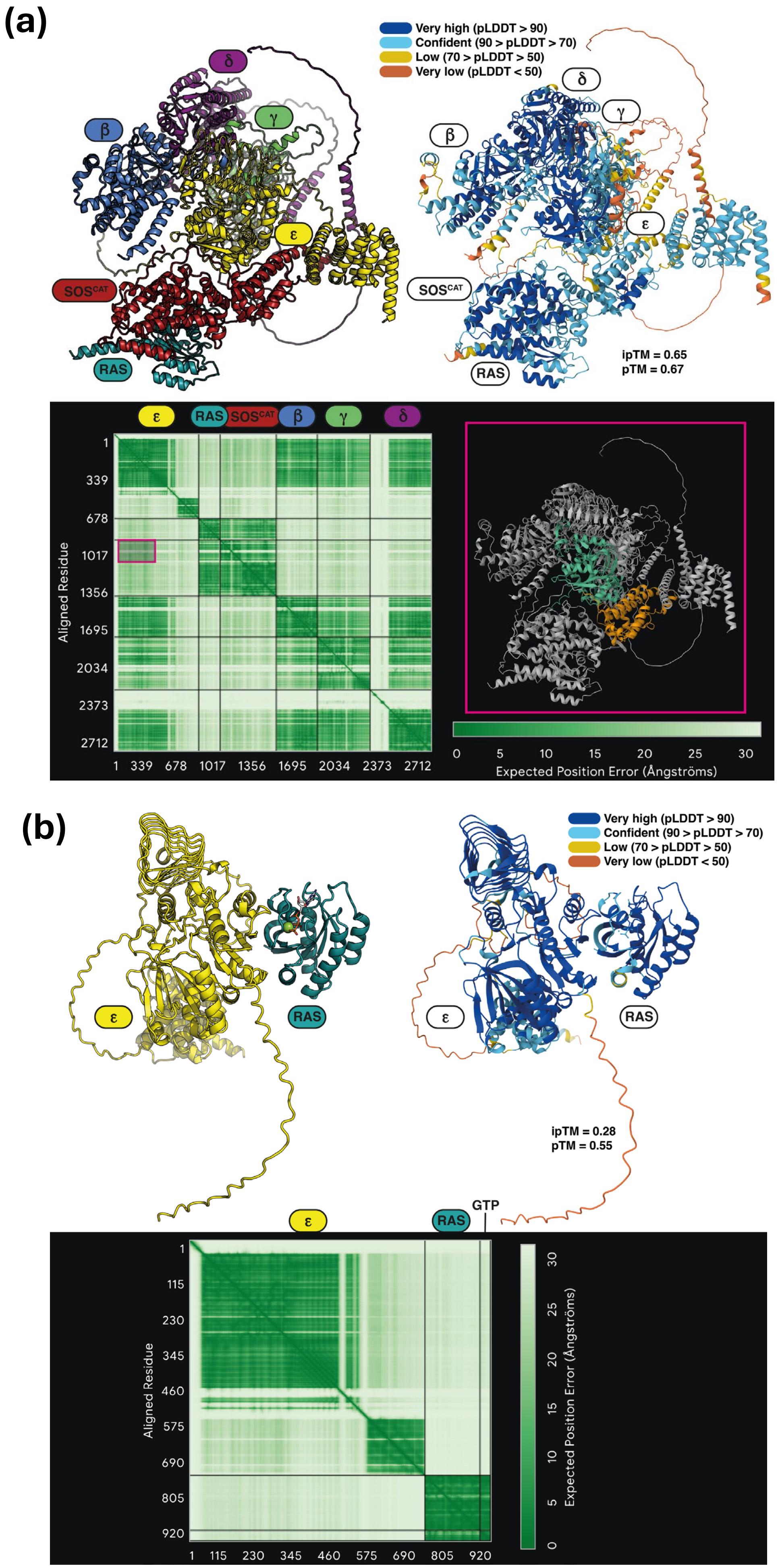


### Suppl Figure 9
